# Environment as a limiting factor of the historical global spread of mungbean

**DOI:** 10.1101/2022.04.27.489711

**Authors:** Pei-Wen Ong, Ya-Ping Lin, Hung-Wei Chen, Cheng-Yu Lo, Marina Burlyaeva, Thomas Noble, Ramakrishnan Nair, Roland Schafleitner, Margarita Vishnyakova, Eric Bishop-von-Wettberg, Maria Samsonova, Sergey Nuzhdin, Chau-Ti Ting, Cheng-Ruei Lee

## Abstract

While the domestication history has been investigated in many crops, the process of cultivation range expansion and factors governing this process received relatively little attention. Here using mungbean (*Vigna radiata* var. *radiata*) as a test case, we investigated the genomes of more than one thousand accessions to illustrate climatic adaptation’s role in dictating the unique routes of cultivation range expansion. Despite the geographical proximity between South and Central Asia, genetic evidence suggests mungbean cultivation first spread from South Asia to Southeast, East, and finally reached Central Asia. Combining evidence from demographic inference, climatic niche modeling, plant morphology, and records from ancient Chinese sources, we showed that the specific route was shaped by the unique combinations of climatic constraints and farmer practices across Asia, which imposed divergent selection favoring higher yield in the south but short-season and more drought-tolerant accessions in the north. Our results suggest that mungbean did not radiate from the domestication center as expected purely under human activity, but instead the spread of mungbean cultivation is highly constrained by climatic adaptation, echoing the idea that human commensals are more difficult to spread through the south-north axis of continents.

## Introduction

Domestication is a process where plants or animals were cultivated by humans, leading to associated genetic and morphological changes. These changes may be intentional from human selection or unintentional as a result of adaptation to the environments of cultivation (Fuller, 2007). Later, the cultivated plants spread out from their initial geographical range (Meyer and Purugganan, 2013), and elucidating the factors affecting the range expansion of crops is another focus of active research (Gutaker et al., 2020). In the old world, during the process of “prehistory food globalization” (Jones et al., 2011), crops originated from distinct regions were transported and grown in Eurasia. Archeological evidence has shown that such “trans-Eurasian exchange” had happened by 1500 BC (Liu et al., 2019), and the proposed spread routes from archeological studies were supported by modern genetic evidence especially in rice (Gutaker et al., 2020) and barley (Lister et al., 2018). Interestingly, the spread may accompany genetic changes for the adaptation to novel environments. For example, in barley, variations in the gene *Photoperiod-H1* (*Ppd-H1*) resulting in the non-responsiveness to longer daylengths were likely associated with the historical expansion to high-latitude regions (Jones et al., 2008; Jones et al., 2016). While these mid-latitude cereals have been extensively studied, investigations of crops originated from other climate zones are needed. Using the South Asian legume mungbean as a test case, here we investigate how climatic adaptation might affect crop spread route and the evolutionary changes making such spread possible.

Mungbean (*Vigna radiata* (L.) Wilczek var. *radiata*), also known as green gram, is an important grain legume in Asia (Nair and Schreinemachers, 2020), providing carbohydrates, protein, folate, and iron for local diets and thereby contributing to food security (Kim et al., 2015). Among pulses, mungbean is capable of tolerating moderate drought or heat stress and has a significant role in rainfed agriculture across arid and semi-arid areas (Pratap et al., 2019), which are likely to have increased vulnerabilities to climate change. Although there have been studies about the genetic diversity of cultivated and wild mungbean (Ha et al., 2021; Kang et al., 2014; Noble et al., 2018; Sangiri et al., 2007), the evolutionary history of cultivated mungbean after domestication still lacks genetic studies. Existing archeological evidence suggests that South Asia is the probable area of mungbean domestication, and at least two independent domestication events have been suggested, including Maharashtra and the eastern Harappan zone (Fuller and Harvey, 2006). The early archeological records suggest that the selection of large seed sizes occurred in the eastern Harappan zone by the 3^rd^ millennium BC and in Maharashtra, dating to the late 2^nd^ to early 1^st^ millennium BC (Fuller and Harvey, 2006). This pulse later spread to mainland Southeast Asia and has been reported in southern Thailand dating to the late first millennium BC (Castillo et al., 2016). Further north, the earliest record of mungbean in China was from the book Qimin Yaoshu (齊民要術, 544 AD). While mungbean is also cultivated in Central Asia today, it was not identified in archaeobotanical evidence ranging from several millennium BC to the medieval period (Miller, 1999; Spengler et al., 2018b; Spengler et al., 2017), suggesting later arrival. While the archaeobotanical studies elucidated the route of mungbean cultivation range expansion, researches are still needed to identify the genetic evidence and factors shaping such spread route.

A recent genetic study revealed that present-day cultivated mungbeans have the same haplotype in the promoter region reducing the expression of *VrMYB26a* (Lin et al., 2022), a candidate gene controlling the important domestication trait, pod shattering, in several *Vigna* species (Takahashi et al., 2020). This suggests the loss of pod shattering phenotype in cultivated mungbean may have a common origin, and despite the archaeobotanical findings of several independent early cultivations of mungbean in South Asia (Fuller and Harvey, 2006), descendants from one of these cultivation origins might have dominated South Asia before the pan-Asia expansion. Since large regions remain archaeologically unexplored, utilization of genetic data can be a crucial complementation to reconstruct crop evolutionary history. Using seed proteins (Tomooka et al., 1992) and isozymes (Dela Vina and Tomooka, 1994), previous studies proposed two expansion routes out of India, one in the south to Southeast Asia and the other in the north along the silk road to China. While later studies used DNA markers to investigate mungbean population structure (Breria et al., 2020; Gwag et al., 2010; Islam and Blair, 2018; Noble et al., 2018; Sandhu and Singh, 2021; Sangiri et al., 2007), few have examined these hypothesized routes in detail. Therefore, genomic examination of the cultivation rage expansion proposed by archaeobotanical studies and the elucidation of its contributing factors are strongly needed.

In this study, we compiled an international effort, reporting a global mungbean diversity panel of more than 1,100 accessions derived from (i) the mungbean mini-core collection of the World Vegetable Centre (WorldVeg) genebank, (ii) the Australian Diversity Panel (ADP), and (iii) the Vavilov Institute (VIR), which hosts a one-century-old collection enriched with mid-latitude Asian accessions that are underrepresented in other genebanks, many of which were old landraces collected by Nikolai I. Vavilov and his teams in the early 20^th^ century (Burlyaeva et al., 2019). These germplasms harbor a wide range of morphological variations (Figure 1A) and constitute the most comprehensive representation of worldwide mungbean genetic variation. We used this resource to investigate the global history of mungbean after domestication to reveal a spread route highly affected by climatic constraints across Asia, eventually shaping the phenotypic characteristics for local adaptation to distinct environments.

**Figure 1.**
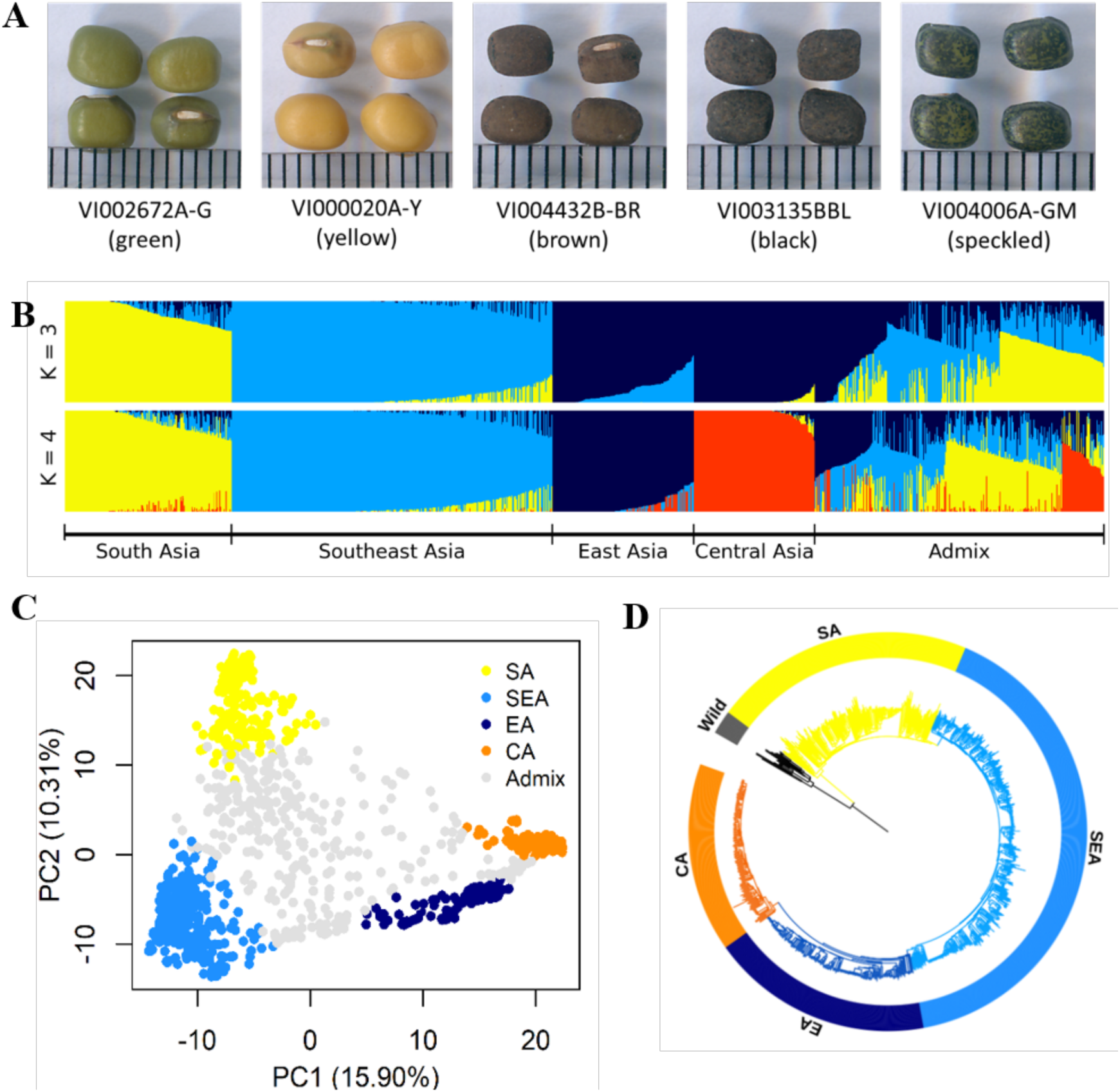
Diversity of worldwide mungbean. (A) Variation in seed colour. (B) ADMIXTURE ancestry coefficients, where accessions were grouped by group assignments (Q ≥ 0.7). (C) Principal component analysis (PCA) plot of 1,092 cultivated mungbean accessions. Accessions were coloured based on their assignment to four inferred genetic groups (Q ≥ 0.7), while accessions with Q < 0.7 were coloured gray. (D) Neighbor-joining (NJ) phylogenetic tree of 788 accessions with Q ≥ 0.7 with wild mungbean as outgroup (black colour).

## Results

### Population structure and spread of mungbean

Using DArTseq, we successfully obtained new genotype data of 290 mungbean accessions from VIR (Supplementary file 1). Together with previous data (Breria et al., 2020; Noble et al., 2018), our final set included 1,108 samples with 16 wild and 1,092 cultivated mungbean. A total of 40,897 single nucleotide polymorphisms (SNPs) were obtained. Of these, 34,469 bi-allelic SNPs, with a missing rate less than 10%, were mapped on 11 chromosomes and retained for subsequent analyses.

The genetic structure was investigated based on the 10,359 LD-pruned SNPs. Principal component analysis (PCA, Figure 1C) showed a triangular pattern of genetic variation among cultivated mungbeans, consistent with previous studies (Breria et al., 2020; Noble et al., 2018; Sokolkova et al., 2020) and ADMIXTURE K = 3 (Figure 1B). The geographic distribution of these genetic groups is not random, as these three groups are distributed in South Asia (India and Pakistan), Southeast Asia (Cambodia, Indonesia, Philippines, Thailand, Vietnam, and Taiwan), and more northernly parts of Asia (China, Korea, Japan, Russia, and Central Asia). As K increased, the cross-validation (CV) error decreased a little after K = 4 (Figure 1-figure supplement 1), where the north group could be further divided (Figure 1B). Therefore, worldwide diversity of cultivated mungbean could be separated into four major genetic groups corresponding to their geography: South Asian (SA), Southeast Asian (SEA), East Asian (EA), and Central Asian (CA) groups. Note that the genetic groups were named after the region where most of their members distribute, and exceptions exist. For example, many EA accessions also distribute in Central Asia, and some SEA accessions were found near the eastern and northeastern coasts of India. Throughout this work, we make clear distinction between genetic group names (e.g., SA) and a geographic region (e.g., South Asia). Therefore, unlike any other previous work in this species, this study incorporates global genetic variation among cultivated mungbean of this important crop.

Using wild progenitor *V. radiata* var. *sublobata* (Wild hereafter) as the outgroup, the accession-level (Figure 1D) and population-level (Figure 2A) phylogenies both suggest CA to be genetically closest to EA. The SEA group is more distant, and SA is the most diverged. This relationship is supported by the outgroup *f*3 tests showing EA and CA share the most genetic drift, followed by SEA and SA (Supplementary file 2). Pairwise *F_ST_* and *d_xy_* also give the same conclusion (Figure 2B). Similarly, the *f*4 tests (Figure 2C) strongly reject the cases where SEA and CA form a clade relative to SA and EA (*f*4(SA,EA;SEA,CA) = 0.016, *Z* = 9.519) or SEA and EA form a clade relative to SA and CA (*f*4(SA,CA;SEA,EA) = 0.021, *Z* = 13.956), again suggesting EA and CA to be closest. With regards to the relationship among Wild, SA, SEA, and EA, *f*4 tests suggest SEA and EA form a clade relative to Wild and SA (non-significant results in *f*4(Wild,SA;EA,SEA) but opposite in other combinations). Notably, both TreeMix (Figure 2A) and the *f*4 test (Figure 2C, *f*4(SA,SEA;CA,EA) = 0.005, *Z* = 6.843) suggest gene flow between SEA and EA. Consistent with archeological evidence of South Asian domestication, the nucleotide diversity (π) decreased from SA (1.0 × 10^−3^) to SEA (7.0 × 10^−4^) and EA (5.0 × 10^−4^), while the CA group has lowest diversity (3.0 × 10^−4^) (Figure 2B). LD also decays the fastest in Wild and then the SA group (Figure 2D), followed by other genetic groups. In summary, all analyses are consistent with our proposed order of cultivated mungbean divergence.

**Figure 2.**
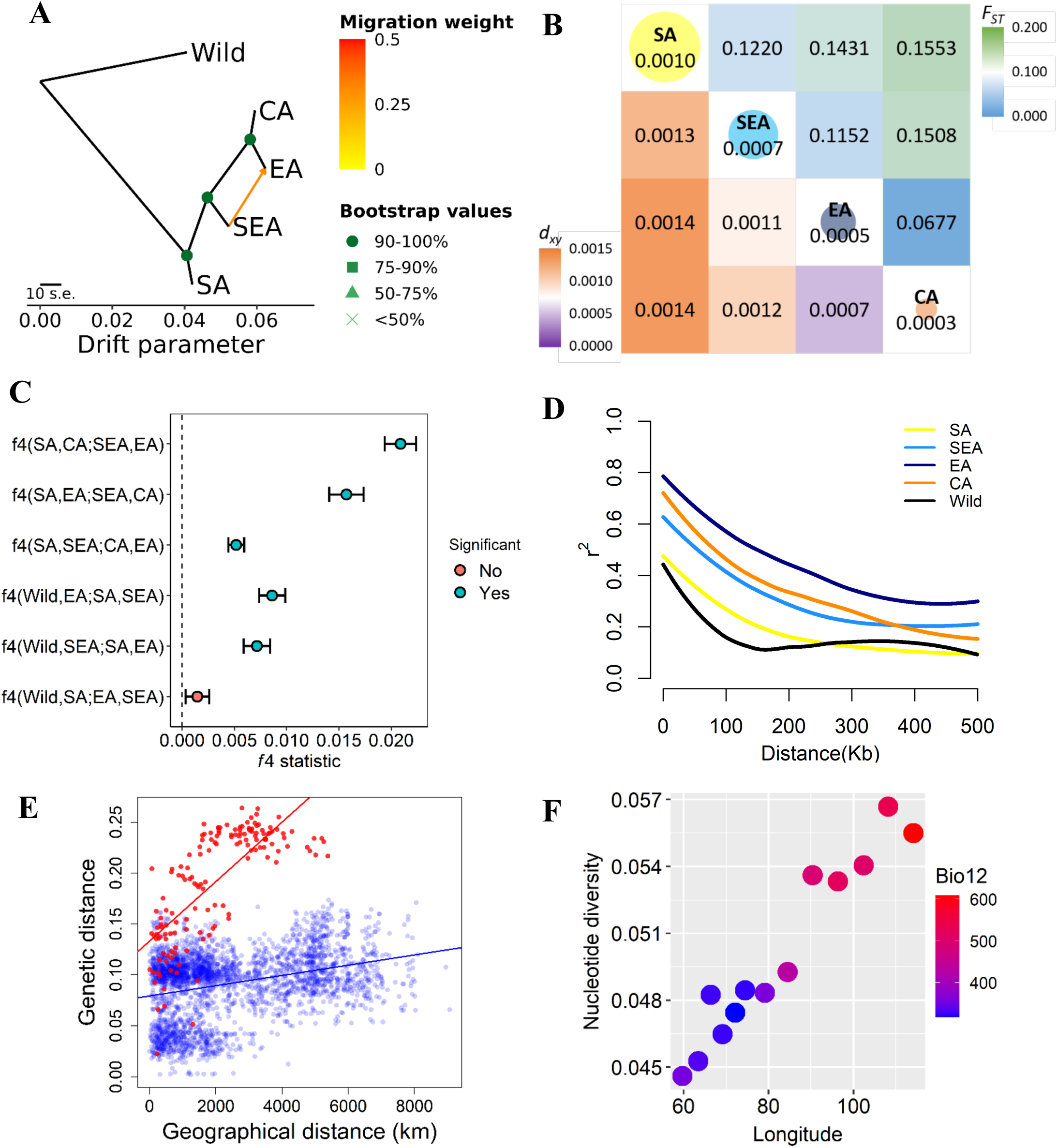
Fine-scale genetic relationship and admixture among four inferred genetic groups. (A) TreeMix topologies with one suggested migration event. Colours on nodes represent support values after 500 bootstraps. (B) Diversity patterns within and between inferred genetic groups as estimated using nucleotide diversity (π in diagonal, where the size of the circle represents the level of π) and population differentiation (*F_ST_* in upper diagonal and *d_xy_* in lower diagonal). (C) *f_4_* statistics. Points represent the mean *f*_4_ statistic and lines are the standard error. Only *f*_4_ statistics with Z-score > |3| are considered statistically significant. The dashed line denotes *f*_4_ = 0. (D) Linkage disequilibrium (LD) decay. (E) Isolation by distance plot of genetic distance versus geographic distance, with the southern group in red circles and the northern group in blue circles. (F) Relationship between Bio12 (annual precipitation) and nucleotide diversity (π) of the EA genetic group across the east-west axis of Asia. Dot colors represent the annual precipitation of each population.

Our proposed demographic history could be confounded by factors such as complex hybridization among groups. For example, SEA and CA might have independently originated from SA and later generated a hybrid population in EA (Figure 2-figure supplement 1A). Other possibilities are that either SEA or CA are the hybrid of other populations (Figure 2-figure supplement 1B and C). We examined these possibilities using *f*3 statistics for all possible trios among the four groups. None of the tests gave a significantly negative *f*3 value (Supplementary file 3), suggesting the lack of a strong alternative model to our proposed relationship among these four groups.

Based on the solid relationship among these genetic groups, we used fastsimcoal2 to model their divergence time, allowing population size change and gene flow at all time points (Figure 2-figure supplement 2A-D). According to this model, after initial domestication, the out-of-India event (when other groups diverged from SA) happened about 8.3 thousand generations ago (kga) with 75% parametric bootstrap range between 4.7 and 11.3 kga. Not until more than five thousand generations later (2.7 kga, 75% range 1.1-4.6 kga) did SEA diverge from the common ancestor of present-day EA and CA. CA diverged from EA only very recently (0.2 kga, 75% range 0.1-0.8 kga). Note that the divergence time was estimated in the number of generations, and the much longer growing seasons in the southern parts of Asia may allow more than one cropping season per year (Mishra et al., 2022; Vir et al., 2016).

Our results suggest the non-South-Asian accessions have a common origin out of India (otherwise these groups would branch off independently from the SA group). Given this, the phylogenetic relationship (Figure 2A) is consistent with the following hypotheses. (1) The east hypothesis: Mungbean expanded eastwards and gave rise to the SEA group. This group might initially occupy northeast South Asia and later expanded to Southeast Asia either through the land or maritime route (Castillo et al., 2016; Fuller et al., 2011). The group later expanded northwards as EA. EA expanded westwards into Central Asia and gave rise to the CA group. (2) The north hypothesis: The group leaving South Asia first entered Central Asia as the EA group. EA expanded eastwards into East Asia through the Inner Asian Mountain Corridor (Stevens et al., 2016). The eastern population of EA expanded southwards as the SEA group, and later the western population of EA diverged as the CA group. (3) The northeast hypothesis: The group leaving South Asia (through either of the above-mentioned routes) was first successfully cultivated in northern East Asia without previously being established in Southeast Asia or Central Asia. The EA group then diverged southwards as SEA and later expanded westwards, giving rise to CA. Consistent with this model, the genetic variation of the EA group gradually declines from east to west, accompanied by the gentlest decline of precipitation per unit geographic distance across Asia (Figure 2F).

While all three hypotheses are consistent with the phylogeny (Figure 2A), the SEA group originated earlier than EA in the east hypothesis but later in the two other hypotheses. The former case predicts higher nucleotide diversity and faster linkage disequilibrium decay in SEA than EA, which is supported by our results (Figure 2B and D). While populations that were established in a region for an extended time could accumulate genetic differentiation, generating patterns of isolation by distance, rapid-spreading populations in newly colonized regions could not (Lee et al., 2017; The 1001 Genomes Consortium, 2016). Using this idea, Mantel’s test revealed a significantly positive correlation between genetic and geographic distances for the SA genetic group (*r* = 0.466, *P* = 0.010), followed by SEA (*r* = 0.252, although not as significant, *P* = 0.069). No such association was found for EA (*r* = 0.030, *P* = 0.142) or CA (*r* = 0.087, *P* = 0.172). In addition, the southern groups (SA and SEA) together (*r* = 0.737, *P* = 0.001) have a much stronger pattern of isolation by distance than the northern groups (EA and CA, *r* = 0.311, *P* = 0.001) (Figure 2E). Using Q ≥ 0.5 instead of Q ≥ 0.7 to assign individuals into genetic groups generated results that are largely consistent (Supplementary file 4). These results are again consistent with the “east hypothesis” that local accessions from the SA and SEA groups were established much earlier than those from EA and CA. Finally, the genetic variation of the EA group is highest in the eastern end and declines westwards (Figure 2F). This does not support the north hypothesis where EA first existed in Central Asia and expanded eastwards.

### Environmental differentiation of the inferred genetic groups

We further examined the possible causes governing the expansion of mungbean cultivation ranges. For a crop to be successfully cultivated in a new environment, dispersal and adaptation are both needed. Being a crop that has lost the ability of pod shattering, the spread of mungbean was governed by commerce or seed exchange. While barriers such as the Himalayas or Hindu Kush may limit human activity, South and Central Asia was already connected by a complex exchange network linking the north of Hindu Kush, Iran, and the Indus Valley as early as about 4 thousand years ago (kya) (Dupuy, 2016; Kohl, 2007; Kohl and Lyonnet, 2008; Lamberg-Karlovsky, 2002; Lombard, 2020; Lyonnet, 2005), and some sites contain diverse crops originated across Asia (Spengler et al., 2021). Similarly, other ancient land or maritime exchange routes existed among South, Southeast, East, and Central Asia (Stevens et al., 2016). This suggests that mungbean could have been transported from South to Central Asia, but our genetic evidence suggests that the present-day CA group did not descend directly from the SA group. Therefore, we investigated whether climatic adaptation, that is, the inability of mungbean to establish in a geographic region after human-mediated long-range expansion, could be a contributing factor.

Multivariate analysis of variance (MANOVA) of eight bioclimatic variables (after removing highly-correlated ones; Supplementary file 5 and 6) indicated strong differentiation in the environmental niche space of the four genetic groups (Supplementary file 7 and 8). PCA of climatic factors clearly reflects geographic structure, where the axis explaining most variation (PC1, 42%) separates north and south groups and is associated with both temperature- and precipitation-related factors (Figure 3A and Supplementary file 9). Consistent with their geographic distribution, overlaps between EA and CA and between SA and SEA were observed. While these analyses were performed using bioclimatic variables from year-round data, we recognized that summer is the cropping season in the north. Parallel analyses using the temperature and precipitation of May, July, and September yielded similar results (Supplementary file 10; Figure 3-figure supplement 1).

**Figure 3.**
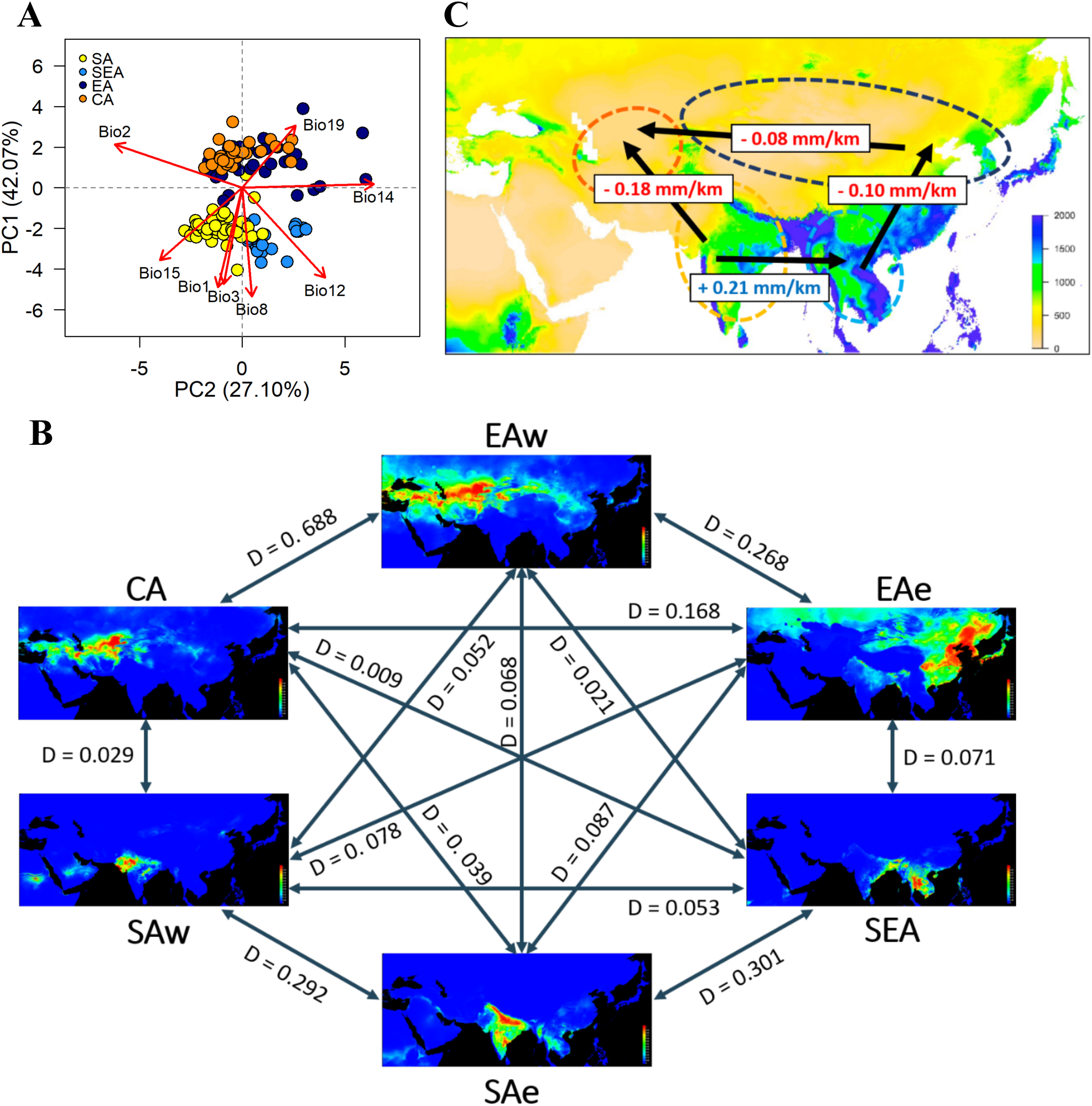
Environmental variation among genetic groups of mungbean. (A) Principal component analysis (PCA) of the eight bioclimatic variables. Samples are coloured according to four inferred genetic groups as indicated in the legend. (B) Predicted distribution at current climate conditions. Red colour indicates high suitability, and blue indicates low suitability. Values between pairs represent niche overlap measured using Schoener’s D, and higher values represent higher overlaps. Abbreviations: SAw: South Asia (west), SAe: South Asia (east); SEA: Southeast Asia; EAe: East Asia (east); EAw: East Asia (west) and CA: Central Asia. (C) Environmental gradient across potential directions of expansion. The value on each arrow indicates a change in annual precipitation per kilometer. The background map is colored according to annual precipitation (Bio12, in mm).

Based on the Köppen climate classification (Köppen, 2011), we categorized the Asian mungbean cultivation range into six major climate zones (Figure 3-figure supplement 2): dry hot (BSh and BWh), dry cold (BSk and BWk), temperate dry summer (Csa), tropical savanna (Aw), Continental (Dwb and Dfb), and temperate wet summer (Cfa and Cwa). The former three are relatively drier than the latter three zones. While SEA and CA are relatively homogeneous, SA and EA have about half of the samples in the dry and non-dry zones (Figure 3-figure supplement 2). We therefore separated SA into SAe and SAw and EA into EAe and EAw, corresponding to the wetter eastern and drier western regions within the SA and EA ranges. Environmental niche modeling revealed distinct suitable regions of these six groups except for CA and EAw, whose geographical ranges largely overlap (Figure 3B). Consistent with PCA, pairwise Schoener’s D values are smallest between the northern and southern groups while largest (suggesting overlaps of niche space) between the eastern and western subsets within north and south (Figure 3B), consistent with PCA that the major axis of climatic difference is between the northern and southern parts of Asia. Analyses using temperature and precipitation from May, July, and September yielded similar results (Figure 3-figure supplement 3). Given a single out-of-India event (Figure 2A), the results suggest it might be easier to first cultivate mungbean in Southeast rather than Central Asia, supporting the east hypothesis.

While both temperature and precipitation variables differ strongly between north and south, one should note that these year-round temperature variables do not correctly reflect conditions in the growing seasons. In the north, mungbean is mostly grown in summer where the temperature is close to the south (Figure 3-figure supplement 4A-C). On the other hand, precipitation differs drastically between the north and south, especially for the CA group, where the summer growing season is the driest of the year (Figure 3-figure supplement 4D). By estimating the regression slope of annual precipitation on geographical distance, we obtained a gradient of precipitation change per unit geographic distance between pairs of genetic groups (Figure 3C). Despite the SA-SEA transect having the steepest gradient (slope = 0.21), the spread from SA to SEA has been accompanied by an increase of precipitation and did not impose drought stress. However, the second highest slope (0.18) is associated with a strong precipitation decrease if the SA group were to disperse to Central Asia. Results from the precipitation of May, July, and September yielded similar conclusion (Figure 3-figure supplement 5). This likely explains why no direct historic spread is observed from South to Central Asia.

### Trait variation among genetic groups

If environmental differences constrained the spread route of mungbean, the current cultivated mungbean accessions occupying distinct environments should have locally adaptive traits for these environments. Indeed, PCA of four trait categories show substantial differences among genetic groups (phenology, reproductive output, and size in field trials, as well as plant weight in lab hydroponic systems, Figure 4A). In the field, CA appears to have the shortest time to flowering, the lowest yield in terms of seed size and pod number, and the smallest leaf size (Figure 4B and Supplementary file 11). On the other hand, SEA accessions maximize seed size, while SA accessions specialize in developing the largest number of pods (Figure 4B). These results suggest that CA has a shorter crop duration, smaller plant size, and less yield, consistent with drought escape phenotypes. This is consistent with the northern short growing season constrained by temperature and daylength (below), as well as the low precipitation during the short season.

**Figure 4.**
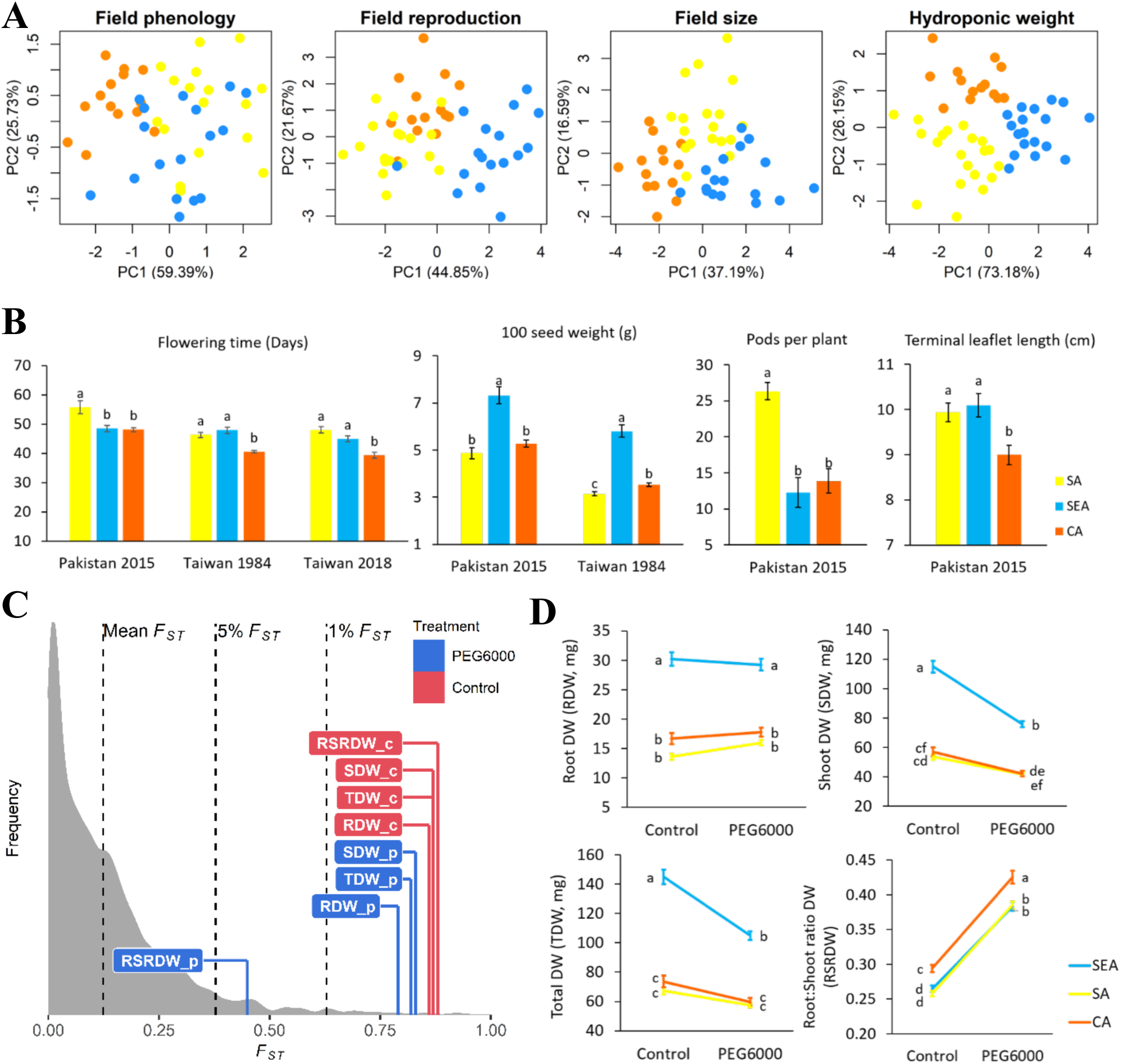
Quantitative trait differentiation among genetic groups. (A) Principal component analysis (PCA) of four trait categories. (B) Trait variability from common gardens in field experiments. (C) Comparison of *Q_ST_*-*F_ST_* for four drought-related traits under two environments. *F_ST_* values (mean, 5% and 1%) were indicated by black dashed lines. The *Q_ST_* for each trait was colored according to treatment and was calculated as Equation 2 in Materials and Methods. Abbreviations: RDW: root dry weight; SDW: shoot dry weight; TDW: total dry weight; RSRDW: root:shoot ratio dry weight; c: control; p: PEG6000. (D) Effect of PEG6000 (−0.6 MPa) on root dry weight (RDW), shoot dry weight (SDW), total dry weight (TDW), and root:shoot ratio dry weight (RSRDW) among genetic groups. Data were expressed as the mean ± standard error. Lowercase letters denote significant differences under Tukey’s HSD in (B) and (D).

In terms of seedling response to drought stress, the *Q_ST_* values of most traits (root, shoot, and whole plant dry weights under control and drought treatments) are higher than the tails of SNP *F_ST_*, suggesting trait evolution driven by divergent selection (Figure 4C; Figure 4-figure supplement 6). Significant treatment, genetic group, and treatment by group interaction effects were observed except on a few occasions (Table 1). Consistent with field observation, SEA has the largest seedling dry weight (Figure 4D). While simulated drought significantly reduced shoot dry weight for all groups, the effect on SEA is especially pronounced (treatment-by-group interaction effect, *F*_2,575_ = 23.55 *P* < 0.001, Table 1 and Figure 4D), consistent with its native habitats with abundant water supply (Figure 3-figure supplement 4D and Supplementary file 12). All groups react to drought in the same way by increasing root:shoot ratio (Figure 4D), suggesting such plastic change may be a strategy to reduce transpiration. Despite the lack of treatment-by-group interaction (*F*_2,575_ = 1.39, *P* > 0.05), CA consistently exhibits a significantly higher root:shoot ratio, a phenotype that is potentially adaptive to its native environment of lower water supply (Figure 3-figure supplement 4D and Supplementary file 12).

**Table 1.** Analysis of variance (ANOVA) F values for the dry weight (mg) of mungbean seedlings across three different genetic groups

| Source of variation | df | RDW | SDW | TDW | RSRDW |
| --- | --- | --- | --- | --- | --- |
| Treatment | 1 | 2.65<sup>n.s.</sup> | 133.26*** | 72.26*** | 978.76*** |
| Genetic group | 2 | 60.63*** | 79.62*** | 76.54*** | 13.27*** |
| Treatment x Genetic group | 2 | 3.29* | 23.55*** | 17.79*** | 1.39<sup>n.s.</sup> |

(df: degrees of freedom; RDW: root dry weight; SDW: shoot dry weight; TDW: total dry weight; RSRDW: root:shoot ratio dry weight. Significance level \* $P < 0.05$ , \*\*\* $P < 0.001$ , n.s. non-significant)

### Support from ancient Chinese sources

Mungbean has been occasionally mentioned in ancient Chinese sources. Here we report the records associated with our proposed mungbean spread route and the underlying mechanisms. The “Classic of Poetry” (Shijing 詩經) contains poems dating between the 11^th^ to 7^th^ centuries BCE near the lower and middle reaches of the Yellow River. While crops (especially soy bean, 菽), vegetables, and many other plants have been mentioned, mungbean was not recorded. This is consistent with our results that mungbean had not reached the northern parts of East Asia at that time (the EA group diverged from the SEA group at around 2.7 kga). The first written record of mungbean in China is in an agricultural encyclopedia Qimin Yaoshu (齊民要術, 544 AD, Chinese text and translation in Supplementary note), whose spatiotemporal background (~1.5 kya near the lower reaches of Yellow River) is again consistent with our estimated origin of the EA group.

Our results suggest that the expansion of the mungbean cultivation range may be associated with the novel phenotypic characteristics potentially adaptive to the new environments. This proposal would be rejected if the novel phenotypic characteristics appeared very recently. In support of our proposal, Xiangshan Yelu (湘山野錄, an essay collection during 1068-1077 AD) recorded that mungbean from the southern parts of Asia had higher yield and larger grains than those in northern China (Chinese text and translation in Supplementary note). Similarly, Tiangong Kaiwu (天工開物, 1637 AD) mentioned that mungbean must be sown during July and August (Chinese text and translation in Supplementary note). The record suggests that the daylength requirement restricts the sowing period of mungbean in the north. Together with the dry summer (Figure 3-figure supplement 4D) and soon-arriving autumn frost, there might be a strong selection favoring accessions with the rapid life cycle. These records suggest the phenotypic characteristics of northern accessions did not originate very recently, and the unique distribution of climatic zones in Asia resulted in not only the specific patterns of expansion but also the evolution of novel phenotypic characteristics in mungbean.

## Discussion

Using mungbean as a test case, we combined population genomics, environmental niche modeling, empirical field and laboratory investigation, and ancient Chinese text analyses to demonstrate the importance of climatic adaptation in dictating the unique patterns of cultivation range expansion after domestication. In this study, we focus on how or when mungbean could be established as part of local agriculture throughout Asia. We showed that after leaving South Asia, mungbean was likely first cultivated in Southeast Asia, East Asia, and finally Central Asia. We acknowledge that our data do not allow us to specify the number of previous out-of-India events that did not leave traces in modern genetic data or their exact routes (for example, whether mungbean expanded from South to Southeast Asia through the land or maritime routes). While there might be multiple attempts to bring mungbean out of India as a commodity for consumption, our results suggest all present-day non-South-Asian accessions have a common out-of-India origin.

### The climate-driven spread route despite historical human activities

Combining archeological records, population genetics, and niche modeling (Figure 2 and 3), our results suggest that after the early cultivation of mungbean in northwestern or southern South Asia (Fuller, 2007; Kingwell-Banham et al., 2015), the large environmental difference may restrict its northwards spread to Central Asia. Mungbean may first spread to eastern South Asia, and the subsequent expansion to Southeast Asia might be facilitated by the environmental similarity between these two regions. This is supported by archaeobotanical remains from the Thai-Malay Peninsula date to ca. 400-100 BCE (Castillo et al., 2016). It took more than five thousand generations until mungbean further spread to northeast Asia, again likely due to the environmental difference. The later appearance of mungbean in northern China is also supported by historical records. After that, the EA group spread across the northern part of Asia within a few thousand generations. Our proposed route suggests that mungbean reached Central Asia at the latest, consistent with its absence from archeological sites in Central Asia, including Turkmenistan and Uzbekistan in the Chalcolithic and Bronze ages (5^th^ to 2^nd^ millennium BC) (Miller, 1999), Southeastern Kazakhstan in the Iron age dating 1^st^ millennium BC (Spengler et al., 2017), and eastern Uzbekistan during the medieval period (800-1100 AD) (Spengler et al., 2018b). In addition, mungbean was only mentioned later by the 18^th^ and early 19^th^ centuries as a pulse grown in the Khiva region of Uzbekistan (Annanepesov and Bababekov, 2003).

In this study, we suggest that the ability to disperse may not be an essential factor restricting mungbean spread from South to Central Asia. Cultivated mungbean has lost the natural ability of pod shattering to disperse seeds, and they mostly traveled through landscapes by human-mediated seed exchange or commerce. Evidence of long-distance human-mediated dispersal of mungbean was available. For example, mungbean seeds have been found near the Red Sea coast of Egypt during the Roman (AD 1–250) period (Van der Veen and Morales, 2015). As early as about four kya, the Bactria–Margiana Archaeological Complex (BMAC) civilization north of the Hindu Kush had extensive contact with the Indus Valley Civilization (Dupuy, 2016; Kohl, 2007; Kohl and Lyonnet, 2008; Lamberg-Karlovsky, 2002; Lombard, 2020; Lyonnet, 2005). By 1,500 BC, the “Trans-Eurasian Exchanges” of major cereal crops has happened (Liu et al., 2019). The frequent crop exchange is evidenced by archaeobotanical findings in the Barikot site (ca. 1200 BC-50 AD) in northern Pakistan (Spengler et al., 2021), where diverse crops were cultivated, including those from West Asia (wheat, barley, pea, and lentil), South Asia (urdbean/mungbean), and likely East Asia (rice). Despite this, in Bronze-age archeological sites north of Hindu Kush, legumes (such as peas and lentils) were observed to a lesser extent than cereals, and South Asian crops were not commonly found (Jeong et al., 2019; Spengler, 2015; Spengler et al., 2014a; Spengler et al., 2018a; Spengler et al., 2014b). Interestingly, archeologists suggested legume’s higher water requirement than cereals may be associated with this pattern, and pea and lentil’s role as winter crops in Southwest Asia may be associated with their earlier appearance in northern Central Asia than other legumes (Spengler et al., 2014a; Spengler et al., 2018a; Spengler et al., 2014b). Therefore, despite the possibility of human-mediated seed dispersal between South and Central Asia, our results and archeological evidence concurred that mungbean arrived in Central Asia at the latest, likely restricted by environmental adaptation.

### Local adaptation of mungbean genetic groups

Despite the profound impact of human-mediated dispersal on the spread of these and many other crops (Herniter et al., 2020; Kistler et al., 2018), in mungbean we suggest adaptation to distinct climatic regimes to be an important factor in the establishment after dispersal. Mungbean is commonly grown under rainfed cultivation and depends on the residual moisture in the fields after the primary crop, thus responding to water stress (Douglas et al., 2020). In the south, a temperature range of 20-30°C and annual precipitation of 600-1,000 mm is optimal for mungbean (Ha and Lee, 2019). In Central Asia, however, the annual precipitation could be as low as 286 mm, greatly below the lower limit required for the southern mungbean. This situation could be further acerbated by the fact that mungbean might not be a highly valued crop under extensive care during cultivation. Indeed, the earliest record of mungbean in China (Qimin Yaoshu 齊民要術, 544 AD) emphasizes its use as green manure. In Central Asia, mungbean is a minor crop (Rani et al., 2018) grown with little input, only in the short duration between successive planting of main crops (which is also the dry season in Central Asia, Supplement file 12 and Figure 3-figure supplement 4) and using residual soil moisture with little irrigation. We suggest that the lack of extensive input subjects mungbean to more substantial local climatic challenges than highly valued high-input crops that receive intensive management, including irrigation. Therefore, the combination of climatic constraints and cultural usage, instead of physical barriers, may have shaped the historical spread route of the mungbean despite extensive human activities across the continent.

In addition to the constraint of soil moisture, other factors may have contributed to the selection of short-season accessions in the north. In the short summer seasons of much of Central Asia, short crop cycling is a requirement. In Uzbekistan, mungbean is often sown in early July after the winter wheat season and harvested before mid-October to avoid delays in the next round of winter wheat and escape frost damage. Therefore, fast-maturing accessions are essential for this production system (Rani et al., 2018). Similar rotation systems using mungbean to restore soil fertility during the short summer season after the harvest of the main crop were also mentioned in ancient Chinese sources (Chen, 1980). Mungbean is a short-day species from the south, and daylength likely limits the window when mungbean could be grown in the north: Chinese texts during the 17^th^ century (Tiangong Kaiwu 天工開物, 1637 AD) specifically mentioned the suitable duration to sow mungbean to control the flowering behavior for maximum yield (Supplementary note). Therefore, unlike in the south where yield appears to be an important selection target, the unique combination of daylength, agricultural practices, soil water availability, and frost damage in the north requires the selection for short-season accessions, likely limiting the direct adoption of southern accessions in the north. Consistent with this, Central Asian accessions have a faster life cycle potentially adaptive to both short growing season and reduced soil water availability, with reduced plant size and lower yield as tradeoffs. These accessions also have increased root:shoot ratio for drought adaptation, similar to findings in rice (Xu et al., 2015), alfalfa (Zhang et al., 2018), and chickpea (Kumar et al., 2012).

About accession sampling and climatic niche modeling, we recognize that not all samples have available spatial data, and we do not have samples from some parts of Asia. For example, while most samples of the SEA group were collected from Taiwan, Thailand, and Philippines, we do not have many samples from the supposed contact zone between SA and SEA (Bangladesh and Myanmar) or between SEA and EA (southern China). If more samples were available from these contact zones, the modeled niche space between SA and SEA and between SEA and EA would be even more similar than the current estimate, strengthening our hypothesis that niche similarity might facilitate the cultivation expansion. On the other hand, clear niche differentiation between SA and CA was evident despite the dense sampling near their contact zone. Based on the Köppen climate classification, South Asia could be roughly separated into two major zones, with the eastern zone slightly more similar to Southeast Asia (Figure 3-figure supplement 2). This partially explained the existence of some SEA accessions in the northeastern coast of India. While the SEA genetic group was named after the geographic region where most of its members were found in the present time, we recognize the possibility that it first occupied northeastern South Asia when it diverged from SA. In that case, the SA-SEA divergence time (4.7 to 11.3 thousand generations ago) might indicate the divergence between the two climate zones within South Asia rather than the expansion of mungbean into Southeast Asia, which may occur much later.

## Conclusion

Our study demonstrates that mungbean’s cultivation range expansion is associated with climatic conditions, which shaped the genetic diversity and contributed to adaptive differentiation among genetic groups. The climatic differences likely also resulted in farmers’ differential emphasis on using it mainly as a grain or green manure crop, further intensifying the phenotypic diversification among regional mungbean accessions that could be used as an invaluable genetic resource for genetic improvement in the future.

## Materials and Methods

### Plant materials and SNP genotyping

A total of 290 cultivated mungbean (*V. radiata* var. *radiata*) accessions were provided by the Vavilov Institute (VIR). Most of the accessions are mainly landraces collected during 1910-1960 are considered these accessions as oldest cultivated mungbean collection from VIR (Burlyaeva et al., 2019). The term landrace, as we use it here, refers to locally adaptive accessions coming from the countries traditionally cultivating them, which also lacks modern genetic improvement. The complete list of materials can be found in Supplementary file 1. Genomic DNA was extracted from a single plant per accession using the QIAGEN Plant Mini DNA kit according to the manufacturer’s instruction with minor modification of pre-warming the AP1 buffer to 65°C and increasing the incubation time of the P3 buffer up to 2 hours on ice to increase DNA yield. DNA samples were sent to Diversity Arrays Technology Pty Ltd, Canberra, Australia for diversity array technology sequence (DArTseq) genotyping.

DArTseq data of 521 accessions from the Australian Diversity Panel (ADP) (Noble et al., 2018) and 297 accessions from the World Vegetable Center (WorldVeg) mini-core (Breria et al., 2020) were also included in this study. In total, our dataset contains more than one thousand accessions (1,092) and covers worldwide diversity of cultivated mungbean representing a wide range of variation in seed colour (Figure 1A). Sixteen wild mungbean (*V. radiata* var. *sublobata*) accessions were included as an outgroup. While all accessions used in this study have the country of origin information, only those from VIR have detailed longitude and latitude information. Therefore, for analyses connecting genetic information and detailed location (the isolation by distance analyses), only the VIR samples were used.

The major goal of this study is to investigate the patterns of population expansion and the underlying ecological causes instead of detailed haplotype analyses of specific genomic regions. For this goal, genomewide SNPs provide similar information as whole-genome sequencing, as have been shown in other species. Compared to other genotyping-by-sequencing technologies, DArTseq has the additional advantage of less missing data among loci or individuals, providing a more robust estimation of population structure.

### SNP calling

Trimmomatic version 0.38 (Bolger et al., 2014) was used to remove adapters based on the manufacturer’s adapter sequences. Reads for each accession were trimmed for low-quality bases with quality scores of Q ≤ 10 using SolexaQA version 3.1.7.1 (Cox et al., 2010) and mapped to the mungbean reference genome (Vradiata_ver6, (Kang et al., 2014) using the Burrows-Wheeler Aligner (BWA) version 0.7.15 (Li and Durbin, 2009). Reads were then sorted and indexed using samtools version 1.4.1 (Li et al., 2009). We used Genome Analysis Toolkit (GATK) version 3.7-0-gcfedb67 (McKenna et al., 2010) to call all sites, including variant and invariant sites. We obtained 1,247,721 sites with a missing rate of < 10% and a minimum quality score of 30. SNP calling was performed using GATK (McKenna et al., 2010). Finally, we used VCFtools version 0.1.13 (Danecek et al., 2011) to remove SNPs with more than two alleles and 10% missing data, resulting in 34,469 filtered SNPs. To reduce non-independence caused by linkage disequilibrium (LD) among SNPs, SNPs were pruned based on a 50-SNP window with a step of 5 SNPs and *r^2^* threshold of 0.5 in PLINK (Purcell et al., 2007). This dataset of 10,359 LD-pruned SNPs (10% missing data) was applied for all analyses related to population genomics unless otherwise noted. For TreeMix that require LD-pruned SNPs with no missing dataset, we used 4,396 LD-pruned SNPs with no missing data.

### Population genetics and differentiation analyses

Population structure was investigated based on 10,359 LD-pruned SNPs using ADMIXTURE (Alexander et al., 2009) with the number of clusters (K) ranging from 1 to 10. The analyses were run ten times for each K value, and cross-validation (CV) error was used to obtain the most probable K value for population structure analysis. ADMIXTURE plots were generated using “Pophelper” in R (Francis, 2017). Genetic groups of accessions were assigned based on ancestry coefficient Q ≥ 0.7, otherwise the accession was considered admixed. The population structure was also examined with principal component analysis (PCA). The neighbor-joining phylogenetic tree was calculated using TASSEL (Trait Analysis by aSSociation, Evolution and Linkage) software version 5.2.60 (Bradbury et al., 2007) and visualized using FigTree version 1.4.4 (http://tree.bio.ed.ac.uk/software/figtree/).

The relationships and gene flow among the four inferred genetic groups were further assessed by TreeMix version 1.12 (Pickrell and Pritchard, 2012) using 4,396 LD-pruned SNPs with no missing data. The analysis was run for 0 to 3 migration events with *V. radiata* var. *sublobata* as an outgroup with a block size of 20 SNPs to account for the effects of LD between SNPs. We estimated one as the optimal number of migration events using the “OptM” in R (Fitak, 2021). Bootstrap support for the resulting observed topology was obtained using 500 bootstrap replicates.

Nucleotide diversity (π) and genetic differentiation (*d_xy_* and *F_ST_*) were estimated in 10 kb windows with pixy version 1.2.7.beta1 (Korunes and Samuk, 2021) using all 1,247,721 invariant and variant sites. LD decay for each genetic group was estimated based on 34,469 non-LD-pruned SNPs using PopLDdecay (Zhang et al., 2019). The curves were fitted by a LOESS function and an LD decay plot was drawn using R.

To investigate the relation among inferred genetic groups, *f*3 and *f*4 statistics were computed based on filtered SNPs using ADMIXTOOLS version 7.0 (Patterson et al., 2012). The *f*3 statistic compares allele frequencies in two populations (A, B) and a target population C. In “outgroup *f*3 statistic”, C is the outgroup and positive values represent the shared genetic drift between A and B. In “admixture *f*3 statistic”, negative values indicate that the C is admixed from A and B. For *f*4 statistics, *f*4(A, B; C, D) measures the shared genetic drift between B populations and C and D after their divergence from outgroup A. A positive value indicates that the B population shares more alleles with D, and a negative value indicates that the B population shares more alleles with C. We used two Mb as a unit of block-jackknife resampling to compute standard errors. The Z-scores with absolute values greater than three are considered statistically significant.

To examine the role of geographic distance in shaping spatial genetic differentiation, Mantel tests with 1,000 permutations were performed for each of the ADMIXTURE-inferred genetic groups (separately for the groups defined by Q ≥ 0.7 or Q ≥ 0.5) using “ade4” in R. Pairwise genetic distance between accessions was estimated based on all sites while the great circle geographic distance was determined using “fields” in R. In addition, the same analysis was conducted for southern and northern groups to examine if there was a south-north pattern of differentiation.

Based on the shape of the phylogenetic tree, we used fastsimcoal2 (Excoffier et al., 2021), which does not rely on whole-genome sequencing, to estimate the split time among genetic groups. Fifty accessions were randomly picked from each genetic group. Population size was allowed to change, and gene flow was allowed among populations. This analysis used all sites covered by the DArT tags (including monomorphic sites), and the mutation rate was set to 1×10^−8^ which was within the range of mutation rates used in eudicots (Barrera-Redondo et al., 2021; Zheng et al., 2022). The models were run using unfolded site frequency spectrum using the major allele in the wild progenitor population (*V*. *radiata* var. *sublobata*) as the ancestral allele. The model was run independently 100 times, each with 100,000 simulations. After obtaining the run with the highest likelihood, we performed parametric bootstrapping 100 times to obtain the 75% confidence intervals of each parameter based on the previous study of Gutaker et al. (2020).

### Ecological niche modelling (ENM)

To understand whether the habitats of genetic groups are differentiated, 248 sampling sites (82 for East Asia, 45 for Southeast Asia, 49 for South Asia and 72 for Central Asia genetic groups), in combination with additional presence records obtained from the Global Biodiversity Information Facility (GBIF, https://www.gbif.org/), were used for the analysis. Using the longitude and latitude information, we extracted the Köppen climate zones (Köppen, 2011) using “kgc” in R (Bryant et al., 2017). After excluding zones with less than five samples, the remaining ten zones were grouped into six categories based on climate similarity: dry hot (BSh and BWh), dry cold (BSk and BWk), temperate dry summer (Csa), tropical savanna (Aw), Continental (Dwb and Dfb), and temperate wet summer (Cfa and Cwa). The former three are relatively dry environments.

Climate layers comprising monthly minimum, maximum, mean temperature, precipitation, and 19 bioclimatic variables were downloaded from the WorldClim database version 1.4 (Hijmans et al., 2005). All climate layers available from WorldClim were created based on climate conditions recorded between 1960 and 1990 at a spatial resolution of 30 arc-seconds (approximately 1 km^2^). To minimize redundancy and model overfitting, pairwise Pearson correlations between the 19 bioclimatic variables were calculated using ENMTools version 1.4.4 (Warren et al., 2010), excluding one of the two variables that has a correlation above 0.8 (Supplementary file 6). As a result, eight bioclimatic variables were used for all further analyses, including Bio1 (annual mean temperature), Bio2 (mean diurnal range), Bio3 (isothermality), Bio8 (mean temperature of wettest quarter), Bio12 (annual precipitation), Bio14 (precipitation of driest month), Bio15 (precipitation seasonality) and Bio19 (precipitation of coldest month). Bioclimatic variables were extracted for each occurrence point using “raster” in R (Hijmans, 2021). PCA and multivariate analysis of variance (MANOVA) were conducted to examine whether there was a significant habitat difference among genetic groups. Ecological niche modelling (ENM) was performed using MAXENT version 3.3.1 (Phillips et al., 2006) to predict the geographic distribution of suitable habitats for cultivated mungbean. The ENM analysis was run with a random seed, a convergence threshold of 5,000 and 10-fold cross-validation. As a measure of the habitat overlaps of the four genetic groups, pairwise of Schoener’s D was calculated using ENMTools. The value ranges from 0 (no niche overlap) to 1 (niche complete overlap). In addition, we carried out the same analyses using monthly temperature and precipitation from May, July, and September.

### Field evaluation

Among the 52 accessions used for laboratory experiments, phenotyping of 49 accessions was conducted at WorldVeg, Taiwan in 1984 and 2018 and at Crop Sciences Institute, National Agricultural Research Centre, Pakistan in 2015. The traits related to phenology (days to 50% flowering), reproduction (100 seed weight, pod length, pods per plant, 1000 seed weight, seeds yield per plant, and seeds per pod), and plant size (petiole length, plant height, plant height at flowering, plant height at maturity, primary leaf length, primary leaf width, terminal leaflet length, and terminal leaflet width) were included. Trait values were inverse normal transformed. The analysis of variance (ANOVA) was performed to test for inferred genetic groups differences for each trait using R software (version 4.1.0).

### Drought phenotyping

A total of 52 accessions with ancestry coefficients Q ≥ 0.7 from three genetic groups (Southeast Asia, South Asia, and Central Asia) were selected for experiments of seedling-stage drought response. The experiment was laid out in a completely randomized design with three replicates of each accession under two treatments (control/drought). The experiment was conducted in two independent batches, and the whole experiment included 624 plants (52 accessions × 2 treatments × 3 plants per treatment × 2 batches).

Mungbean seeds were surface-sterilized with 10% bleach for 10 mins and rinsed with distilled water for three times. Seeds were treated with 70% ethanol for 5 mins and washed three times in distilled water. The sterilized seeds were germinated on wet filter paper in petri dishes for 3 days. The experiment was conducted in a 740FLED-2D plant growth chamber (HiPoint, Taiwan) at a temperature of 25 ± 1°C and 12 hours of photoperiod (light ratios of red: green: blue 3: 1: 1) with light intensity 350 µmol m^−2^s^−1^ and relative humidity at 60 ± 5%. The seedlings were then transplanted to a hydroponic system with half-strength Hoagland nutrient solution (Phytotechnology Laboratory, USA) and were grown for six days before drought stress started. The nutrient solution was changed on alternate days and the pH of the solution was adjusted to 6.0 with 1M KOH or 1M HCl.

For drought treatment, seedlings of mungbean were exposed to polyethylene glycol (PEG)-induced drought stress for five days. The solution of PEG6000 with an osmotic potential of −0.6 MPa was prepared by adding PEG6000 (Sigma-Aldrich, Germany) to the nutrient solution according to Michel and Kaufmann (1973), and pH was also adjusted to 6.0. The seedlings grown with the nutrient solution under the same environmental conditions were considered as controls.

At the end of the experiment, plants were evaluated for shoot dry weight (SDW) and root dry weight (RDW), measured on digital balance after oven-drying at 70°C for 48 hours. All traits were analysed by mixed-model ANOVA with the treatment (control/drought) and the genetic group as fixed effects. The models included accessions as a random effect nested within genetic groups and a random effect of batches. Tukey’s test was conducted to compare genetic groups. All statistics were performed using JMP v3.0.0 (SAS Institute, 2016).

### *Q_ST_*-*F_ST_* comparisons

For each trait, quantitative trait divergence (*Q_ST_*) was calculated separately with respect to each treatment. Our root and shoot weight experiment used a selfed-progeny design, using the self-fertilized seeds from each accession as replicates, as recommended for partially inbred species (Goudet and Buchi, 2006). For the selfed-progeny design of inbred species, (Equation 1) *Q_ST_* = V_B_ / (V_B_ + V_Fam_), where V_B_ is the among-population variance component and V_Fam_ is the within-population among-family variance component (Goudet and Buchi, 2006). Variance components were estimated using a model with genetic groups, accessions nested within genetic groups, and batches as random factors. To accommodate the possibility that mungbean is not completely selfing, we also applied (Equation 2) *Q_ST_* = (1 + *f*) V_B_ / ((1 + *f*) V_B_ + 2V_AW_) (Goudet and Buchi, 2006), where *f* is the inbreeding coefficient (estimated by VCFtools as 0.8425), V_B_ is the among-population variance component, and V_AW_ is the additive genetic variance within genetic groups estimated by the kinship matrix using TASSEL software (Bradbury et al., 2007). The results and conclusions are similar to our previous version. The *F_ST_* was calculated only using accessions in the phenotyping experiment.

## Acknowledgements

We thank Chia-Yu Chen and Shang-Ying Tien for their assistance in sample preparation. C-RL was funded by 107-2923-B-002-004-MY3 and 110-2628-B-002-027 from the Ministry of Science and Technology, Taiwan. C-TT was funded by 107-2923-B-002-004-MY3 from the Ministry of Science and Technology, Taiwan. Y-PL was supported by 110-2313-B-125-001-MY3 from the Ministry of Science and Technology, Taiwan. RN and RS were funded by the Australian Center for International Agricultural Research (ACIAR) through the projects on International Mungbean Improvement Network (CIM-2014-079 and CROP-2019-144) and by the strategic long-term donors to the World Vegetable Center: Republic of China (Taiwan), UK aid from the UK government, United States Agency for International Development (USAID), Germany, Thailand, Philippines, Korea, and Japan. EBvW was supported by USDA Multistate Hatch NE1710. EBvW and MS were supported by Russian Scientific Fund Project No. 18-46-08001 and the Ministry of Science and Higher Education of the Russian Federation as part of World-class Research Center program: Advanced Digital Technologies (contract No. 075-15-2020-934 dated 17.11.2020). SN was supported by the Zumberge foundation.

## Data Availability

Sequences generated in this study are available under NCBI BioProject PRJN809503. Accession names, GPS coordinates, and NCBI accession numbers of the Vavilov Institute accessions are available under Supplementary file 1. Plant trait data are available at Dryad https://doi.org/10.5061/dryad.d7wm37q3h. Sequences and accession information of the World Vegetable Centre mini-core and the Australian Diversity Panel collections were obtained from Breria et al. (2020) and Noble et al. (2018).

## Supplementary note

Text analysis and translation of ancient Chinese texts regarding mungbean

Qimin Yaoshu (齊民要術, about 544 AD)

Qimin Yaoshu, compiled by Sixie Jia (賈思勰), is one of the earliest and most complete agricultural sources in China, detailing agricultural techniques near the lower reaches of Yellow River at that era. This is the earliest record of mungbean in China, demonstrating mungbean has reached northern China at that time and is consistent with our estimates of population divergence time. The popularity of mungbean is demonstrated by it being mentioned multiple times under different contexts, most notably as a green manure:

「若糞不可得者,五六月中,穊種菉豆,至七月、八月,犁掩殺之。如以糞糞田, 則良美與糞不殊,又省功力。」

Translation: “Should feces be unavailable, during May and June one could grow mungbean. Until July or August, one could plow mungbean plants into the soil. This is equivalent to using feces to manure the land. This is as good as using feces and saves efforts.”

Notice that the months used in ancient China are slightly different from the Gregorian calendar.

Xiangshan Yelu (湘山野錄, 1068-1077 AD)

Xiangshan Yelu was written by a monk, Wen-Ying (文瑩), recording anecdotes during that era. Its records about the Emperor Zhenzong of Song (宋真宗, 968-1022 AD) detailed the phenotypes of Indian mungbean at that time:

「真宗深念稼穡,聞占城稻耐旱、西天綠豆子多而粒大,各遣使以珍貨求其種。占 城得種二十石,至今在處播之。西天中印土得菉豆種二石,不知今之菉豆是否?」

Translation: “Zhenzong of Song deeply concerned about agriculture. He heard Champa rice being drought tolerant and mungbean from India produce numerous and large seeds. Diplomats were sent to exchange the seeds with treasure. Twenty dans of Champa rice were obtained and propagated everywhere. Two dans of mungbean were obtained from India, but it is unclear whether the mungbean today descended from these.”

“Dan” (石) is a unit of volume in ancient China and is called “Koku” in Japanese. The exact amount varied with time.

The texts provide us with two pieces of important information. First, mungbean from South Asia (likely also includes the SEA genetic groups if accessions near eastern India and Bangladesh were included) at that time had higher yield and larger seeds than native mungbean accessions in northern China, consistent with our results on trait divergence. Second, compared to the clear success of Champa rice in China, it was unclear whether those southern accessions had prospered in northern China, likely suggesting an unsuccessful introduction of southern high-yield and large-seeded accessions to the north.

Tiangong Kaiwu (天工開物, 1637 AD)

Tiangong Kaiwu is a famous Chinese encyclopedia compiled by Song Yingxing (宋應 星). While it mostly covers technologies at that time, a section about agricultural practices covers mungbean:

「綠豆必小暑方種,未及小暑而種,則其苗蔓延數尺,結莢甚稀。若過期至於處 暑,則隨時開花結莢,顆粒亦少。」

Translation: “Mungbean must be sown at or after Xiaoshu (Gregorian 7-8 July). Being sown before Xiaoshu, mungbean stems would spread for meters with few pods set. Being sown as late as Chushu (Gregorian 23-24 August), the plants would flower and set pods at any time, also with low yield.”

As a short-day plant, being sown too early when the days are too long, mungbean would have mostly vegetative growth. Being sown too late when the days are too short, flowering would be induced too quickly before sufficient vegetative development. In addition to our results that short-season accessions were favored in the north due to the requirement for drought escape, this source provides us with another support that mungbean could only be sown in a narrow time window due to daylength requirement. Given the autumn frost damage in the north, not being able to be sown earlier restricts the growing season length in the north, limiting the adoption of southern long-season accessions.

---

**Figure 1-figure supplement 1.**
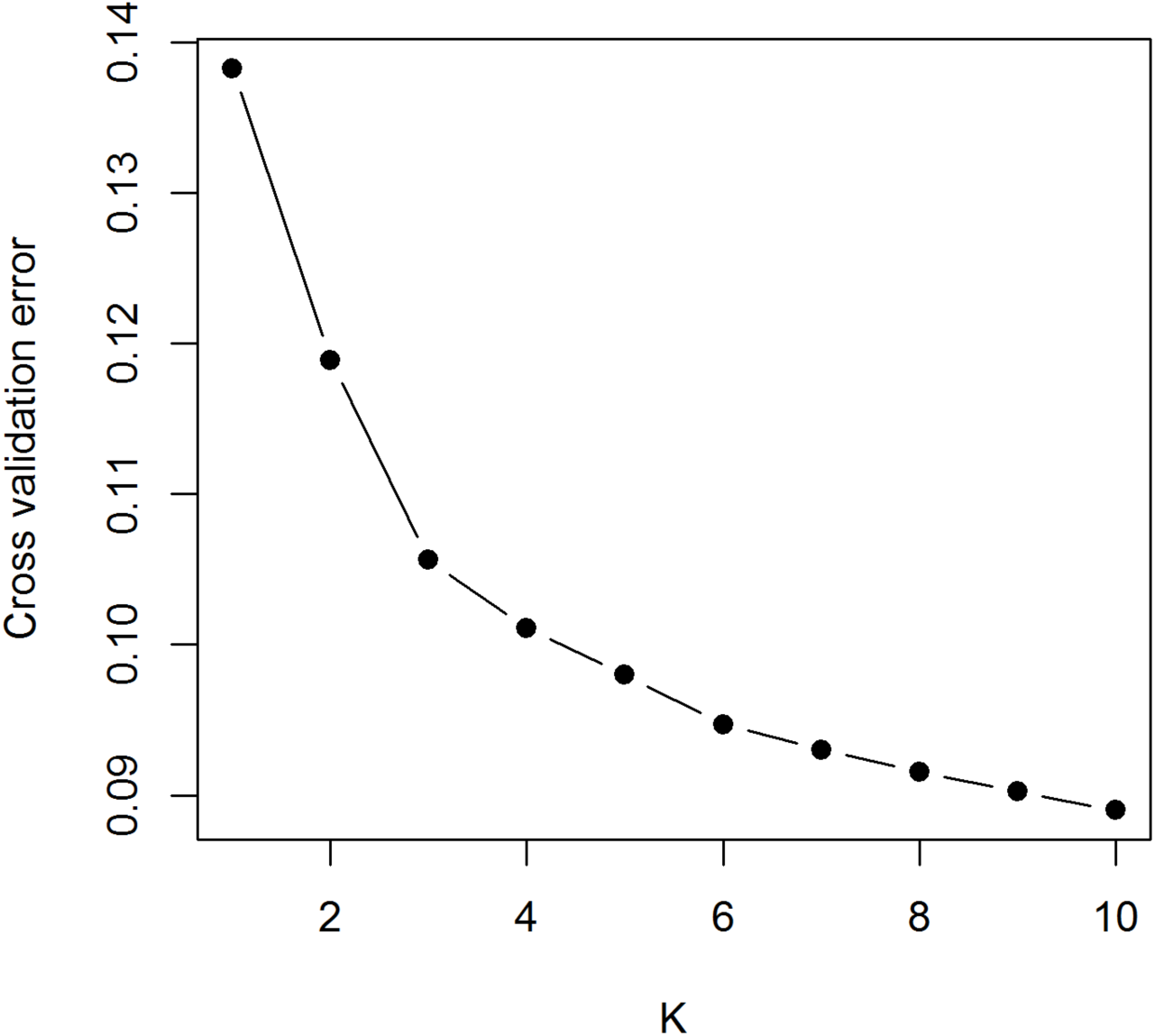
Cross-validation (CV) errors of ADMIXTURE. Means of CV errors were calculated based on K values ranging 1 to 10 with 10 independent runs.

**Figure 2-figure supplement 1.**
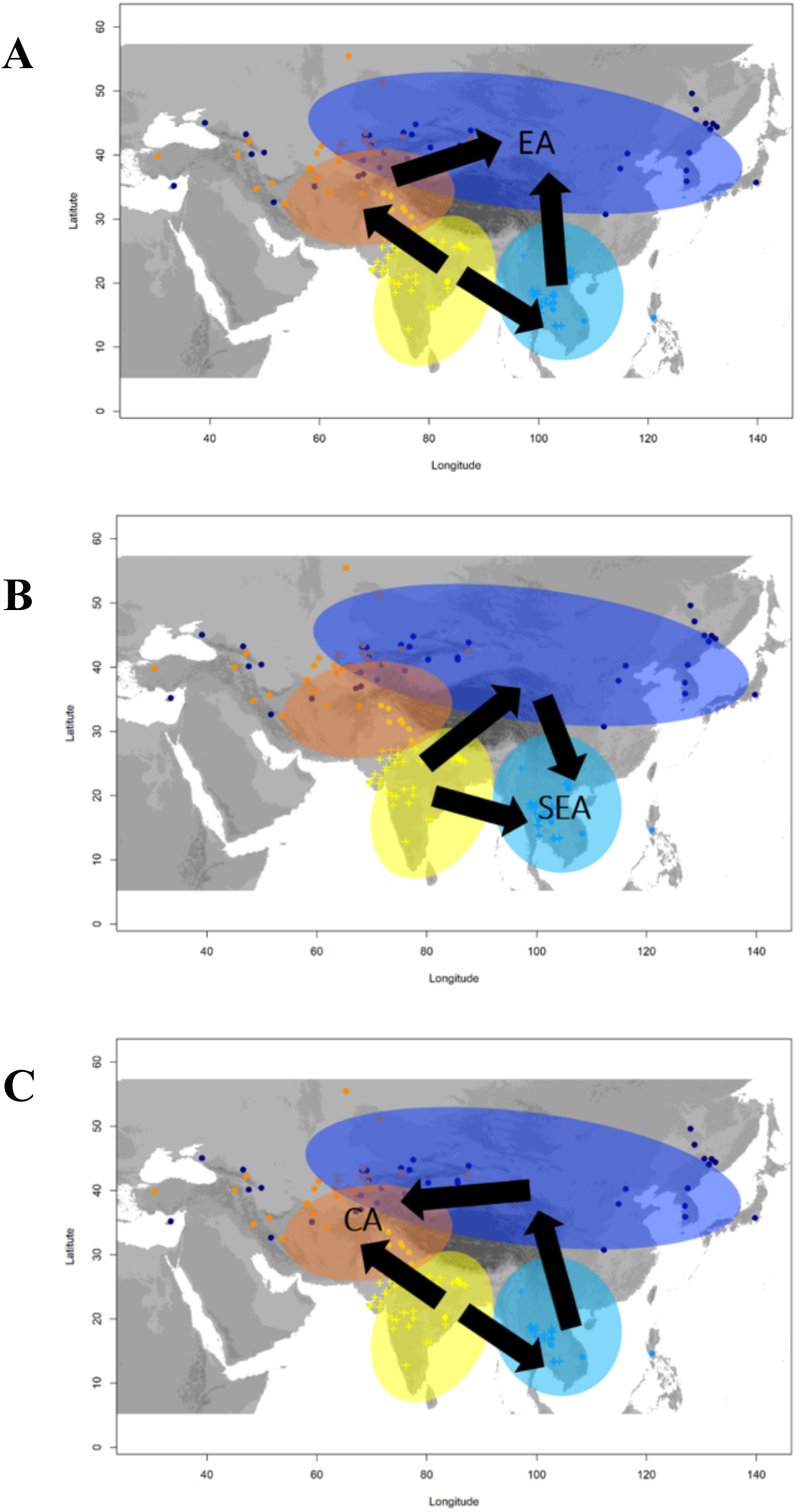
Schematic representation to investigate presence of admixture in a target population from two source populations using admixture f3-statistics. (A) f3(EA; SEA, CA), (B) f3(SEA; SA, EA) and (C) f3(CA; EA, SA). Coloured circles indicate the geographic area occupied by distinct genetic groups. Arrows indicates the possible direction of expansion and admixture among populations.

**Figure 2-figure supplement 2.**
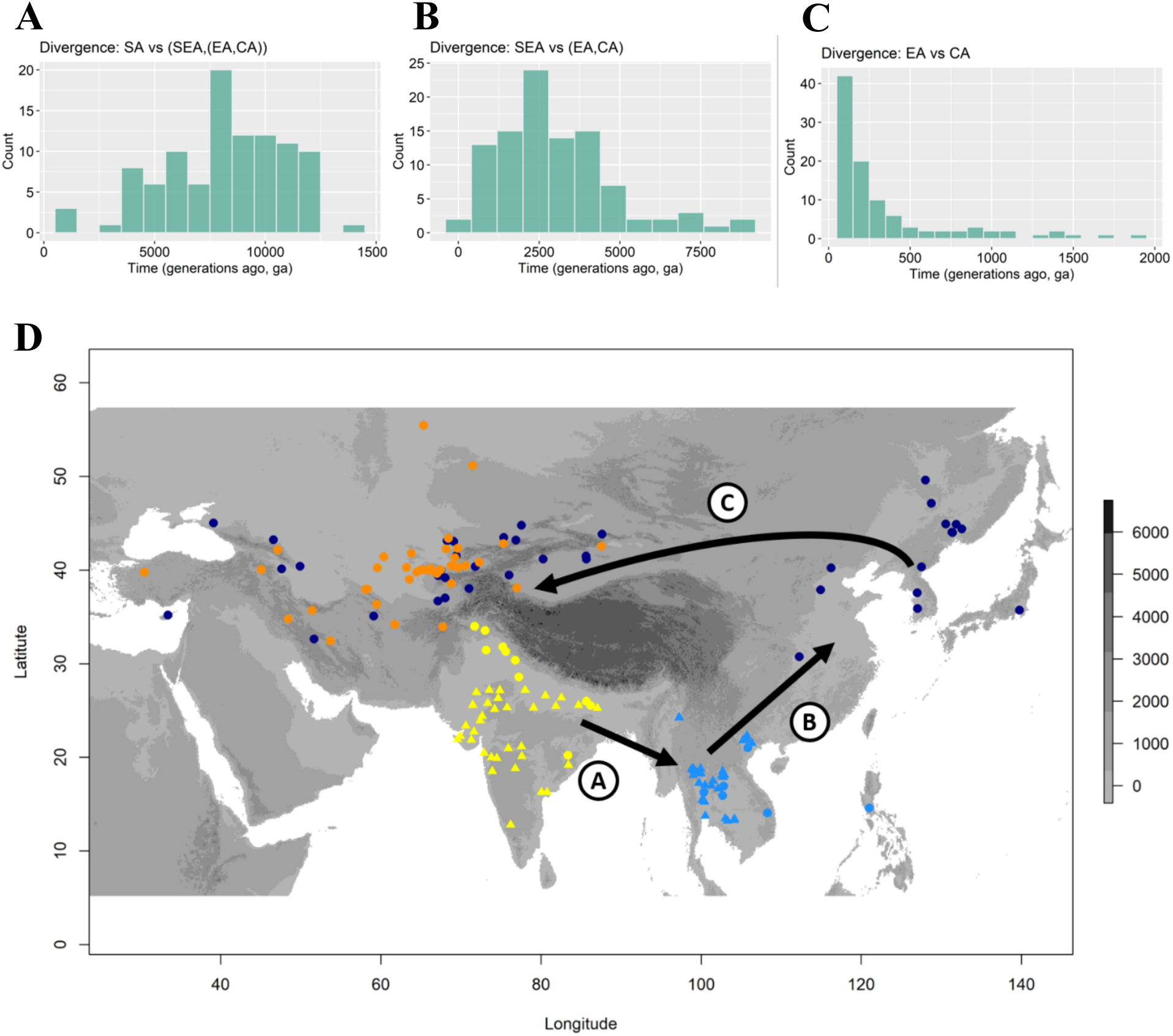
Estimates of divergence time and inferred mungbean movement over time across Asia. The histograms of the divergence times represent (A) split time between SA and (SEA,(EA,CA)), (B) split time between SEA and (EA,CA), and (C) split time between EA and CA. (D) Geographic distribution of mungbean accessions and proposed mungbean spread routes. Exact locations for VIR accessions (filled circle) and GBIF records (filled triangle) are provided. Each accession was coloured the same as an inferred genetic group using ADMIXTURE in Figure 1. Arrow indicates the possible expansion directions. The map was shaded as a gray colour representing altitude (meters above sea level).

**Figure 3-figure supplement 1.**
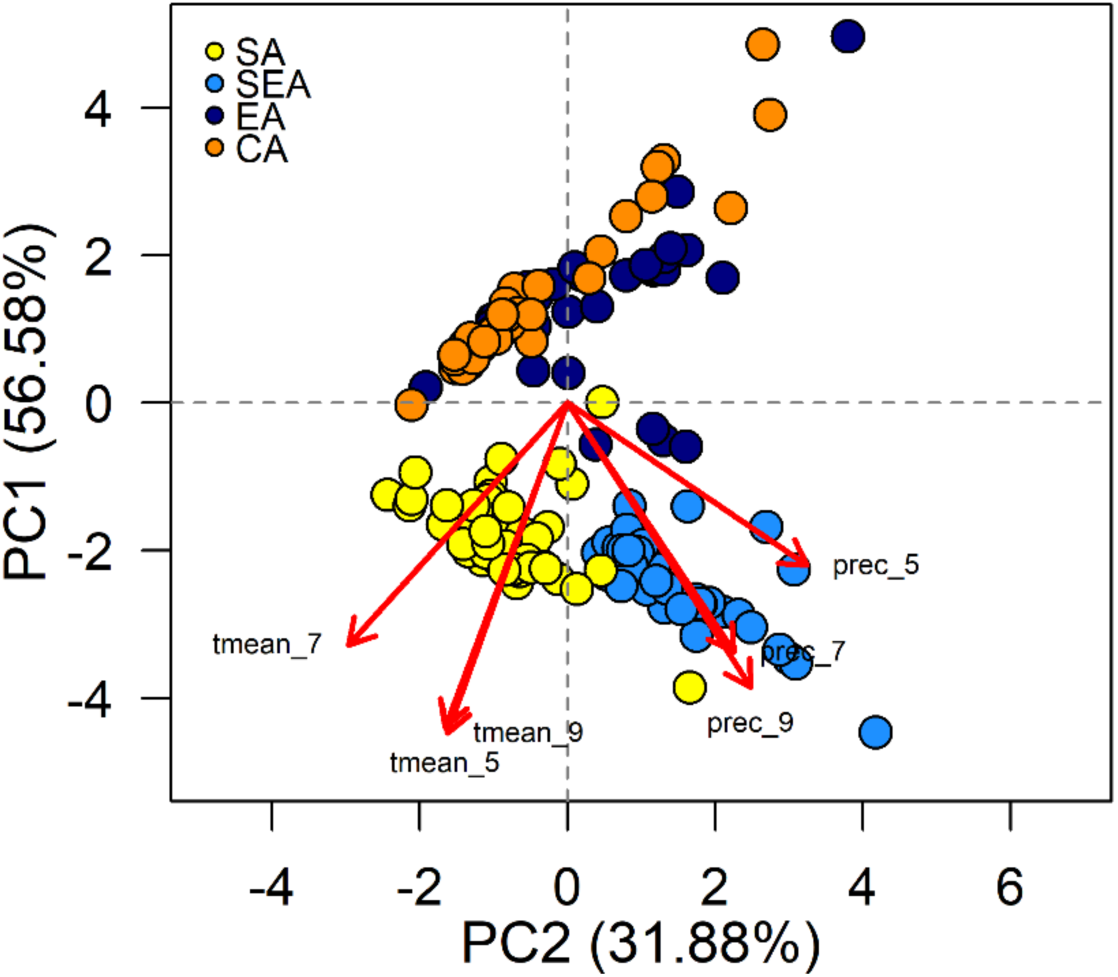
Principal component analysis (PCA) of the growing season climatic data including temperature and precipitation of May, July and September. Samples are coloured according to four inferred genetic groups, as indicated in the legend.

**Figure 3-figure supplement 2.**
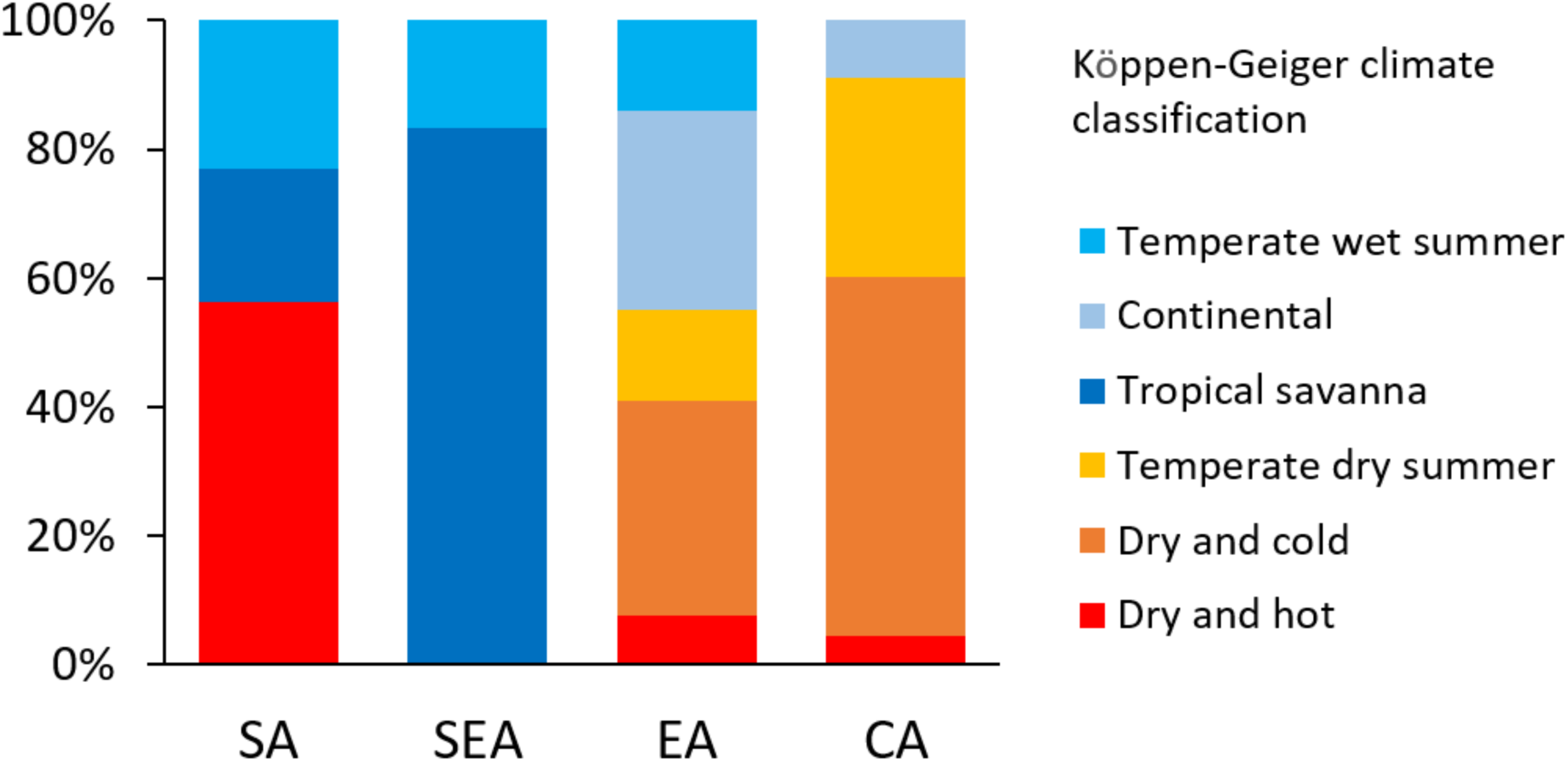
The distribution of accessions in major climate zones according to the Köppen climate classification (Köppen, 2011).

**Figure 3-figure supplement 3.**
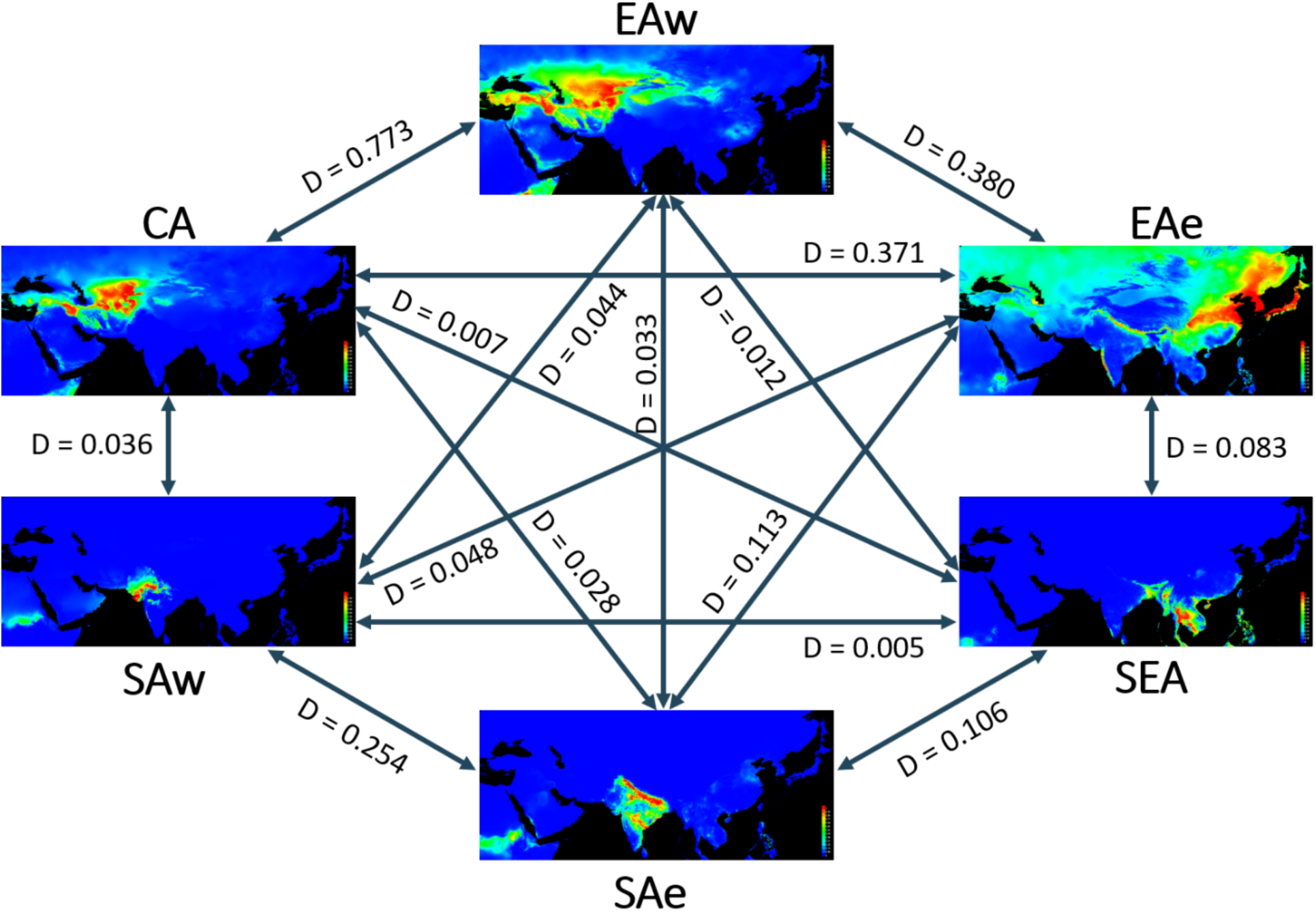
Predicted distributions of six groups based on monthly temperature and precipitation (May, July and September) during the summer growing season. Red colour indicates high suitability and blue indicates low suitability. Values between groups represent niche overlap measured using Schoener’s D. Abbreviations: SAw: South Asia (west), SAe: South Asia (east); SEA: Southeast Asia; EAe: East Asia (east); EAw: East Asia (west) and CA: Central Asia.

**Figure 3-figure supplement 4.**
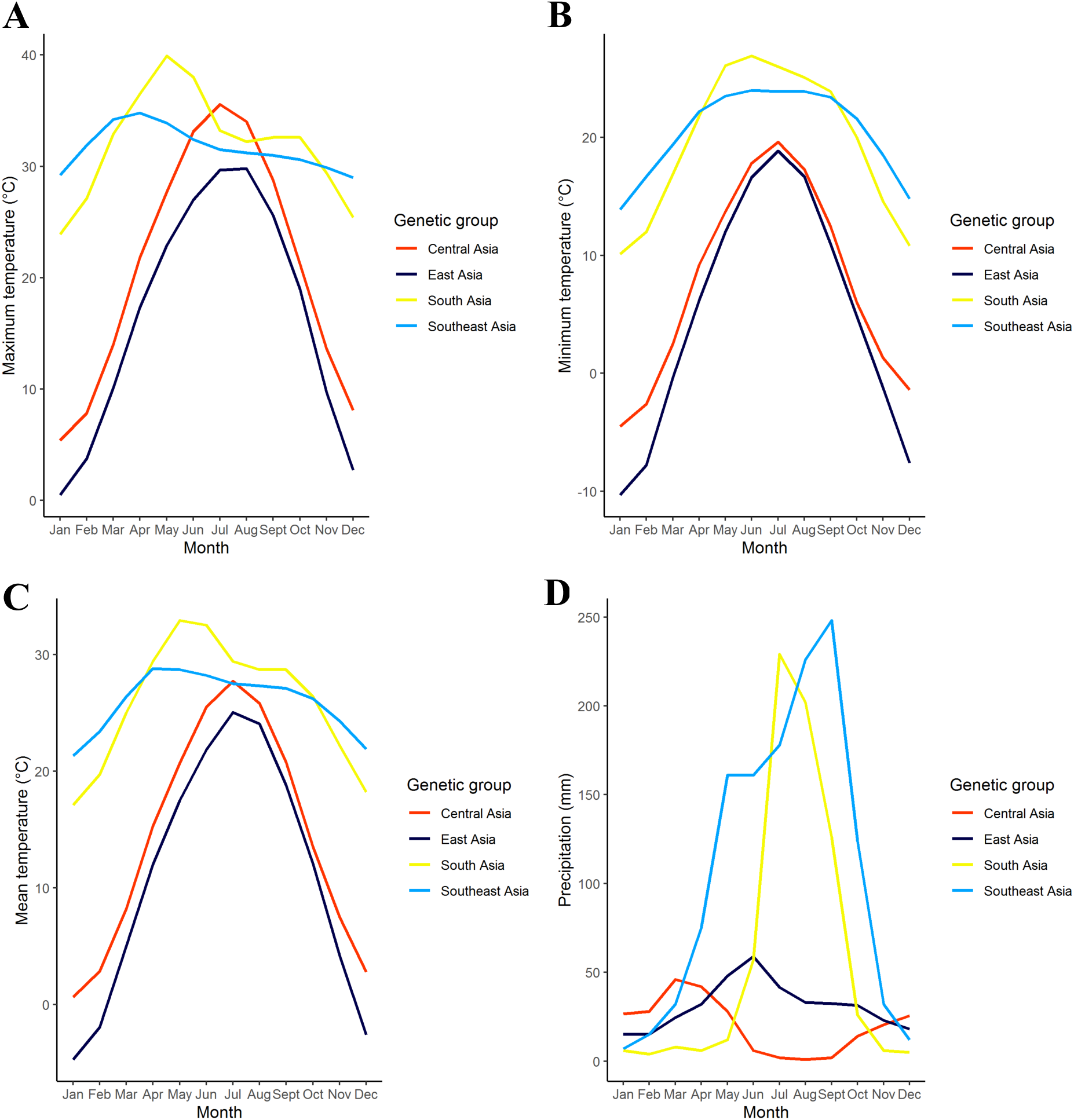
Monthly temperature and precipitation variations among the four genetic groups. Monthly (A) maximum temperature, (B) minimum temperature, (C) mean temperature and (D) precipitation were computed based on median value among all accessions of a group. Genetic group were coloured the same as in Figure. 1.

**Figure 3-figure supplement 5.**
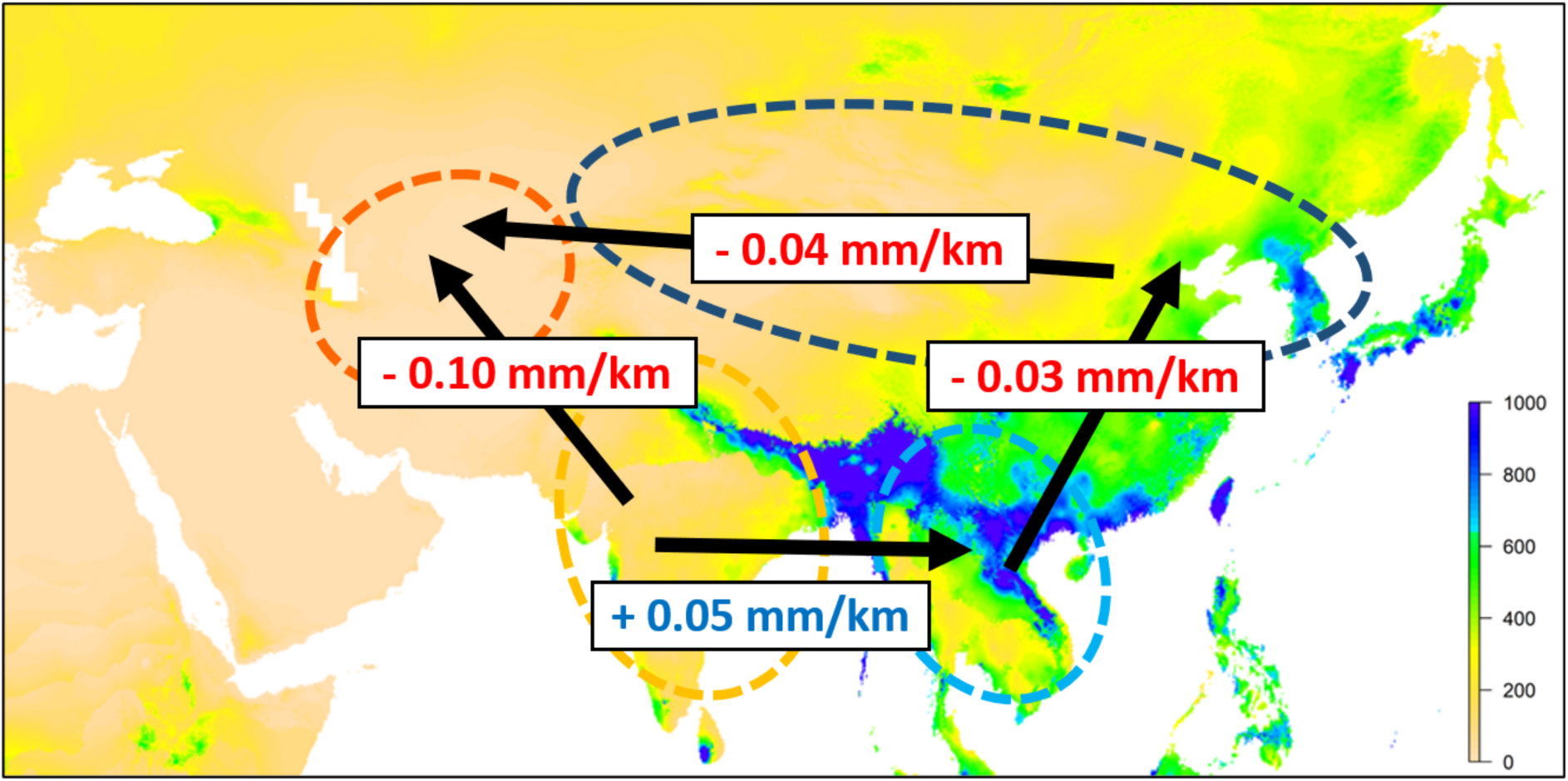
Environmental gradient across Asia. The value on each arrow indicates a change in mean precipitation for May, July and September (growth season) per kilometer. The background map is colored according to summer precipitation (Bio18, precipitation of warmest quarter, in mm).

**Figure 3-figure supplement 6.**
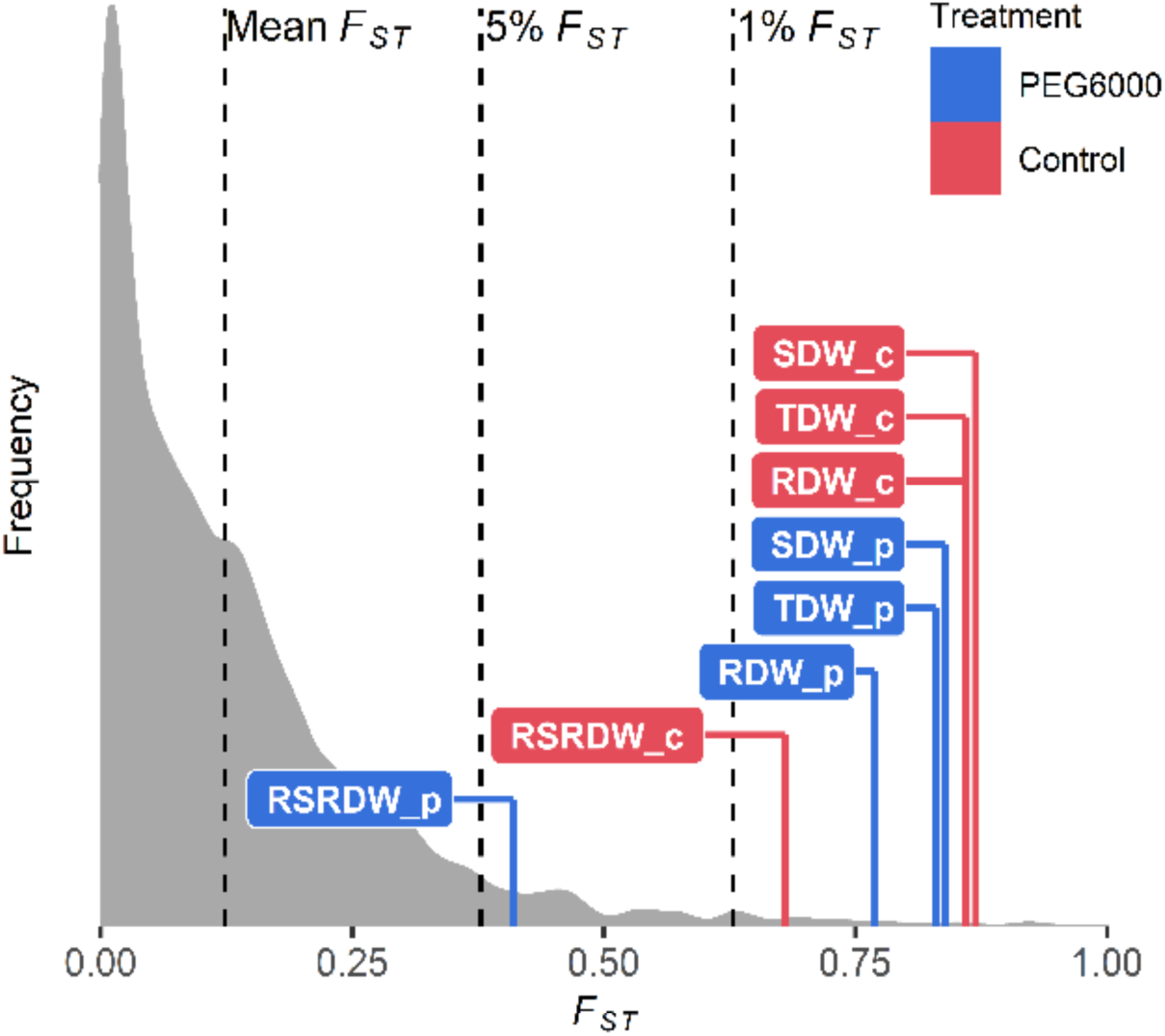
Comparison of *Q_ST_*-*F_ST_* for four drought-related traits under two environments. *F_ST_* values (mean, 5% and 1%) were indicated by black dashed lines. The *Q_ST_* for each trait was colored according to treatment and was calculated as Equation 1 in Materials and Methods. Abbreviations: RDW: root dry weight; SDW: shoot dry weight; TDW: total dry weight; RSRDW: root:shoot ratio dry weight; c: control; p: PEG6000.

**Supplementary file 1.** Mungbean accessions from Vavilov Institute (VIR) collection

| Sample name | Country | Latitude | Longitude | NCBI SRA accession | NCBI Biosample accession |
| --- | --- | --- | --- | --- | --- |
| 1.B | USA | NA | NA | SRR18125483 | SAMN26179197 |
| 100.A | Cyprus | 35.1856 | 33.3823 | SRR18125482 | SAMN26179198 |
| 101.A | Morocco | 33.9716 | -6.8498 | SRR18125266 | SAMN26179199 |
| 102.A | Israel | 32.4971 | 35.4973 | SRR18125200 | SAMN26179200 |
| 103.A | Indonesia | -6.5971 | 106.806 | SRR18125367 | SAMN26179201 |
| 104.B | Russia | 44.9003 | 131.8351 | SRR18125356 | SAMN26179202 |
| 105.A | Chile | -36.8305 | -73.1167 | SRR18125345 | SAMN26179203 |
| 106.A | India | 22.5587 | 88.2911 | SRR18125298 | SAMN26179204 |
| 107.A | India | 22.5726 | 88.3639 | SRR18125287 | SAMN26179205 |
| 108.A | India | 25.9821 | 85.6486 | SRR18125276 | SAMN26179206 |
| 109.A | Italy | 41.9028 | 12.4964 | SRR18125481 | SAMN26179207 |
| 10A | China | 47.1216 | 128.7382 | SRR18125398 | SAMN26179208 |
| 110.B | Portugal | 38.7223 | -9.1393 | SRR18125387 | SAMN26179209 |
| 111.A | Portugal | 38.7223 | -9.1393 | SRR18125376 | SAMN26179210 |
| 112.A | Russia | 45.0347 | 39.0978 | SRR18125329 | SAMN26179211 |
| 113.A | Tajikistan | 40.2675 | 69.6453 | SRR18125318 | SAMN26179212 |
| 114.B | India | 31.5204 | 74.3587 | SRR18125307 | SAMN26179213 |
| 115.B | India | 31.5204 | 74.3587 | SRR18125476 | SAMN26179214 |
| 116.A | Russia | 49.6152 | 127.9945 | SRR18125465 | SAMN26179215 |
| 116.B | Russia | 49.6152 | 127.9945 | SRR18125454 | SAMN26179216 |
| 117.B | Russia | 44.0281 | 131.3273 | SRR18125264 | SAMN26179217 |
| 118.A | Russia | 44.0118 | 131.3835 | SRR18125253 | SAMN26179218 |
| 119.A | Russia | 44.39 | 132.558 | SRR18125242 | SAMN26179219 |
| 11A | China | 47.1216 | 128.7382 | SRR18125231 | SAMN26179220 |
| 120.A | Argentina | -32.8895 | -68.8458 | SRR18125434 | SAMN26179221 |
| 122.A | Japan | 31.5969 | 130.5571 | SRR18125423 | SAMN26179222 |
| 123.B | Kyrgyzstan | 55.4649 | 65.3054 | SRR18125412 | SAMN26179223 |
| 124.B | Ethiopia | 12.9545 | 36.1573 | SRR18125223 | SAMN26179224 |
| 125.A | China | 42.5246 | 87.5396 | SRR18125212 | SAMN26179225 |
| 126.A | China | 42.5246 | 87.5396 | SRR18125201 | SAMN26179226 |
| 127.A | China | 42.5246 | 87.5396 | SRR18125199 | SAMN26179227 |
| 128.A | Democratic Republic of the Congo | 1.9293 | 30.0492 | SRR18125198 | SAMN26179228 |
| 128.B | Democratic Republic of the Congo | 1.9293 | 30.0492 | SRR18125197 | SAMN26179229 |
| 129.A | Korea | 37.5665 | 126.978 | SRR18125196 | SAMN26179230 |
| 12A | China | 47.1216 | 128.7382 | SRR18125195 | SAMN26179231 |
| 13.A | Iran | 32.4279 | 53.688 | SRR18125194 | SAMN26179232 |
| 130.A | Korea | 37.5665 | 126.978 | SRR18125371 | SAMN26179233 |
| 131.A | Korea | 37.5665 | 126.978 | SRR18125370 | SAMN26179234 |
| 132.B | Korea | 40.3399 | 127.5101 | SRR18125369 | SAMN26179235 |
| 133.A | Korea | 40.3399 | 127.5101 | SRR18125368 | SAMN26179236 |
| 134.A | Korea | 40.3399 | 127.5101 | SRR18125366 | SAMN26179237 |
| 135.A | Korea | 40.3399 | 127.5101 | SRR18125365 | SAMN26179238 |
| 136.A | Korea | 40.3399 | 127.5101 | SRR18125364 | SAMN26179239 |
| 137.A | China | 42.5246 | 87.5396 | SRR18125363 | SAMN26179240 |
| 138.A | China | 39.4677 | 75.9938 | SRR18125362 | SAMN26179241 |
| 139.A | Uzbekistan | 41.2995 | 69.2401 | SRR18125361 | SAMN26179242 |
| 14.B | Uzbekistan | 39.7681 | 64.4556 | SRR18125360 | SAMN26179243 |
| 140.B | China | 42.5246 | 87.5396 | SRR18125359 | SAMN26179244 |
| 141.A | China | 43.8256 | 87.6168 | SRR18125358 | SAMN26179245 |
| 142.B | China | 42.5246 | 87.5396 | SRR18125357 | SAMN26179246 |
| 144.A | Ukraine | 48.4647 | 35.0462 | SRR18125355 | SAMN26179247 |
| 145.A | Brazil | -22.9329 | -47.0738 | SRR18125354 | SAMN26179248 |
| 146.B | Kazakhstan | 43.222 | 76.8512 | SRR18125353 | SAMN26179249 |
| 147.A | Turkey | 39.7646 | 30.4559 | SRR18125352 | SAMN26179250 |
| 148.B | Turkey | 39.7646 | 30.4559 | SRR18125351 | SAMN26179251 |
| 149.A | Ukraine | 48.0386 | 30.9497 | SRR18125350 | SAMN26179252 |
| 151.A | Tajikistan | 38.5598 | 68.787 | SRR18125349 | SAMN26179253 |
| 154.A | Brazil | -14.235 | -51.9253 | SRR18125348 | SAMN26179254 |
| 155.A | United Kingdom | 52.3555 | -1.1743 | SRR18125347 | SAMN26179255 |
| 156.A | India | 25.9821 | 85.6486 | SRR18125346 | SAMN26179256 |
| 157.A | India | 25.9821 | 85.6486 | SRR18125344 | SAMN26179257 |
| 158.A | India | 25.9821 | 85.6486 | SRR18125343 | SAMN26179258 |
| 159.B | India | 25.9821 | 85.6486 | SRR18125342 | SAMN26179259 |
| 16.A | Uzbekistan | 39.7681 | 64.4556 | SRR18125341 | SAMN26179260 |
| 160.A | Uzbekistan | 40.8154 | 72.2837 | SRR18125340 | SAMN26179261 |
| 161.A | Uzbekistan | 40.8154 | 72.2837 | SRR18125339 | SAMN26179262 |
| 162.A | South Africa | -26.7145 | 27.097 | SRR18125338 | SAMN26179263 |
| 163.A | South Africa | -26.7145 | 27.097 | SRR18125337 | SAMN26179264 |
| 164.A | South Africa | -26.7145 | 27.097 | SRR18125336 | SAMN26179265 |
| 165.B | USA | 39.9526 | -75.1652 | SRR18125299 | SAMN26179266 |
| 166.A | USA | 39.9526 | -75.1652 | SRR18125297 | SAMN26179267 |
| 167.B | Kyrgyzstan | 42.8224 | 75.3179 | SRR18125296 | SAMN26179268 |
| 168.A | Kyrgyzstan | 42.8224 | 75.3179 | SRR18125295 | SAMN26179269 |
| 17.B | Uzbekistan | 39.7681 | 64.4556 | SRR18125294 | SAMN26179270 |
| 170.A | Russia | 43.2562 | 46.5893 | SRR18125293 | SAMN26179271 |
| 171.A | Russia | 43.2562 | 46.5893 | SRR18125292 | SAMN26179272 |
| 172.B | Iran | 32.4279 | 53.688 | SRR18125291 | SAMN26179273 |
| 173.A | Iran | 32.4279 | 53.688 | SRR18125290 | SAMN26179274 |
| 174.A | Korea | 40.3399 | 127.5101 | SRR18125289 | SAMN26179275 |
| 175.A | Uzbekistan | 40.2504 | 63.2032 | SRR18125288 | SAMN26179276 |
| 175.B | Uzbekistan | 40.2504 | 63.2032 | SRR18125286 | SAMN26179277 |
| 176.A | Uzbekistan | 40.2504 | 63.2032 | SRR18125285 | SAMN26179278 |
| 177.A | Uzbekistan | 40.2504 | 63.2032 | SRR18125284 | SAMN26179279 |
| 178.A | Uzbekistan | 40.2504 | 63.2032 | SRR18125283 | SAMN26179280 |
| 179.A | Uzbekistan | 39.9208 | 66.4271 | SRR18125282 | SAMN26179281 |
| 18.B | Uzbekistan | 39.7681 | 64.4556 | SRR18125281 | SAMN26179282 |
| 180.A | Uzbekistan | 39.9208 | 66.4271 | SRR18125280 | SAMN26179283 |
| 181.A | Uzbekistan | 39.9208 | 66.4271 | SRR18125279 | SAMN26179284 |
| 182.B | Uzbekistan | 39.9208 | 66.4271 | SRR18125278 | SAMN26179285 |
| 183.A | Uzbekistan | 40.3734 | 71.7978 | SRR18125277 | SAMN26179286 |
| 184.A | China | 41.482754 | 85.626702 | SRR18125275 | SAMN26179287 |
| 187.A | China | 38.10222 | 76.993816 | SRR18125274 | SAMN26179288 |
| 188.B | China | 41.1675 | 80.2634 | SRR18125273 | SAMN26179289 |
| 189.A | China | 41.1675 | 80.2634 | SRR18125272 | SAMN26179290 |
| 19.B | Iran | 32.4279 | 53.688 | SRR18125271 | SAMN26179291 |
| 190.A | China | 42.9513 | 89.1898 | SRR18125270 | SAMN26179292 |
| 191.A | China | 41.175324 | 85.660861 | SRR18125269 | SAMN26179293 |
| 192.A | India | 31.8183 | 75.2071 | SRR18125268 | SAMN26179294 |
| 193.A | India | 31.326 | 75.5762 | SRR18125267 | SAMN26179295 |
| 193.B | India | 31.326 | 75.5762 | SRR18125265 | SAMN26179296 |
| 194.A | India | 30.3752 | 76.7821 | SRR18125480 | SAMN26179297 |
| 195.A | India | 28.7041 | 77.1025 | SRR18125407 | SAMN26179298 |
| 195.B | India | 28.7041 | 77.1025 | SRR18125406 | SAMN26179299 |
| 197.A | India | 17.6599 | 75.9064 | SRR18125405 | SAMN26179300 |
| 199.A | Senegal | 14.4974 | -14.4524 | SRR18125404 | SAMN26179301 |
| 1A | USA | NA | NA | SRR18125403 | SAMN26179302 |
| 201.B | Pakistan | 31.5204 | 74.3587 | SRR18125402 | SAMN26179303 |
| 202.B | Canada | 43.6502 | -79.9036 | SRR18125401 | SAMN26179304 |
| 203.A | Ethiopia | 9.3126 | 42.1227 | SRR18125400 | SAMN26179305 |
| 204.B | Indonesia | -6.5971 | 106.806 | SRR18125399 | SAMN26179306 |
| 205.A | Indonesia | -6.5971 | 106.806 | SRR18125397 | SAMN26179307 |
| 205.B | Indonesia | -6.5971 | 106.806 | SRR18125396 | SAMN26179308 |
| 206.A | Hungary | 47.1625 | 19.5033 | SRR18125395 | SAMN26179309 |
| 207.A | Viet Nam | 14.0583 | 108.2772 | SRR18125394 | SAMN26179310 |
| 21.B | Kazakhstan | 51.1605 | 71.4704 | SRR18125393 | SAMN26179311 |
| 210.A | Afghanistan | 36.6926 | 67.118 | SRR18125392 | SAMN26179312 |
| 212.A | Viet Nam | 21.0278 | 105.8342 | SRR18125391 | SAMN26179313 |
| 215.A | Guinea | 8.5383 | -9.4728 | SRR18125390 | SAMN26179314 |
| 216.A | Uzbekistan | 39.972132 | 65.558096 | SRR18125389 | SAMN26179315 |
| 217.B | Uzbekistan | 40.162885 | 66.227209 | SRR18125388 | SAMN26179316 |
| 218.A | Uzbekistan | 40.013465 | 64.943243 | SRR18125386 | SAMN26179317 |
| 219.A | Turkmenistan | 40.243331 | 59.540314 | SRR18125385 | SAMN26179318 |
| 22.B | Kazakhstan | 42.3417 | 69.5901 | SRR18125384 | SAMN26179319 |
| 220.A | Turkmenistan | 40.243331 | 59.540314 | SRR18125383 | SAMN26179320 |
| 223.A | Uzbekistan | 40.483568 | 70.546311 | SRR18125382 | SAMN26179321 |
| 224.A | Iran | 32.6539 | 51.666 | SRR18125381 | SAMN26179322 |
| 225.A | Pakistan | 30.3753 | 69.3451 | SRR18125380 | SAMN26179323 |
| 226.B | Pakistan | 30.3753 | 69.3451 | SRR18125379 | SAMN26179324 |
| 227.A | Pakistan | 30.3753 | 69.3451 | SRR18125378 | SAMN26179325 |
| 228.B | Turkmenistan | 39.0041 | 63.5688 | SRR18125377 | SAMN26179326 |
| 229.A | Pakistan | 31.4504 | 73.135 | SRR18125375 | SAMN26179327 |
| 23.A | Kazakhstan | 42.3417 | 69.5901 | SRR18125374 | SAMN26179328 |
| 230.A | Pakistan | 31.4504 | 73.135 | SRR18125373 | SAMN26179329 |
| 230.B | Pakistan | 31.4504 | 73.135 | SRR18125372 | SAMN26179330 |
| 231.A | Tanzania | -6.369 | 34.8888 | SRR18125335 | SAMN26179331 |
| 232.A | Iran | 34.3277 | 47.0778 | SRR18125334 | SAMN26179332 |
| 233.A | Iran | 32.4279 | 53.688 | SRR18125333 | SAMN26179333 |
| 234.A | India | 28.6139 | 77.209 | SRR18125332 | SAMN26179334 |
| 235.A | Pakistan | 33.5651 | 73.0169 | SRR18125331 | SAMN26179335 |
| 235.B | Pakistan | 33.5651 | 73.0169 | SRR18125330 | SAMN26179336 |
| 236.A | Pakistan | 34.0155 | 71.6888 | SRR18125328 | SAMN26179337 |
| 237.A | Pakistan | 28.6001 | 77.227 | SRR18125327 | SAMN26179338 |
| 238.A | Egypt | 26.8206 | 30.8025 | SRR18125326 | SAMN26179339 |
| 239.A | Kazakhstan | 44.7689 | 77.5573 | SRR18125325 | SAMN26179340 |
| 24.A | Uzbekistan | 41.4065 | 60.3685 | SRR18125324 | SAMN26179341 |
| 240.A | Uzbekistan | 39.4065 | 67.1418 | SRR18125323 | SAMN26179342 |
| 240.B | Uzbekistan | 39.4065 | 67.1418 | SRR18125322 | SAMN26179343 |
| 241.A | Uzbekistan | 40.023044 | 67.433724 | SRR18125321 | SAMN26179344 |
| 242.A | Uzbekistan | 39.982851 | 67.486778 | SRR18125320 | SAMN26179345 |
| 244.A | Uzbekistan | 41.773406 | 63.780613 | SRR18125319 | SAMN26179346 |
| 245.A | Kazakhstan | 43.47491 | 75.335144 | SRR18125317 | SAMN26179347 |
| 246.A | Algeria | 35.6971 | -0.6308 | SRR18125316 | SAMN26179348 |
| 247.A | Tanzania | -5.0425 | 32.8197 | SRR18125315 | SAMN26179349 |
| 249.A | Australia | -32.9283 | 151.7817 | SRR18125314 | SAMN26179350 |
| 25.A | Uzbekistan | 41.4065 | 60.3685 | SRR18125313 | SAMN26179351 |
| 250.B | Russia | NA | NA | SRR18125312 | SAMN26179352 |
| 251.B | USA | NA | NA | SRR18125311 | SAMN26179353 |
| 252.A | Afghanistan | 34.7602 | 69.8121 | SRR18125310 | SAMN26179354 |
| 253.A | Afghanistan | 34.7602 | 69.8121 | SRR18125309 | SAMN26179355 |
| 254.A | Yemen | 15.5527 | 48.5164 | SRR18125308 | SAMN26179356 |
| 254.B | Yemen | 15.5527 | 48.5164 | SRR18125306 | SAMN26179357 |
| 255.A | Afghanistan | 36.6926 | 67.118 | SRR18125305 | SAMN26179358 |
| 256.A | Afghanistan | 36.6926 | 67.118 | SRR18125304 | SAMN26179359 |
| 257.B | Afghanistan | 36.6926 | 67.118 | SRR18125303 | SAMN26179360 |
| 258.B | Afghanistan | 33.9391 | 67.71 | SRR18125302 | SAMN26179361 |
| 259.B | Colombia | 4.5709 | -74.2973 | SRR18125301 | SAMN26179362 |
| 26.B | Uzbekistan | 41.4065 | 60.3685 | SRR18125300 | SAMN26179363 |
| 260.A | Philippines | 14.5995 | 120.9842 | SRR18125479 | SAMN26179364 |
| 261.B | Kenya | -1.2921 | 36.8219 | SRR18125478 | SAMN26179365 |
| 262.A | Kenya | -1.2921 | 36.8219 | SRR18125477 | SAMN26179366 |
| 263.A | Kenya | -1.2921 | 36.8219 | SRR18125475 | SAMN26179367 |
| 264.A | Kenya | -1.2921 | 36.8219 | SRR18125474 | SAMN26179368 |
| 265.B | Kenya | -1.2921 | 36.8219 | SRR18125473 | SAMN26179369 |
| 266.A | Kenya | -1.2921 | 36.8219 | SRR18125472 | SAMN26179370 |
| 267.A | Kenya | -1.2921 | 36.8219 | SRR18125471 | SAMN26179371 |
| 268.A | Kenya | -1.2921 | 36.8219 | SRR18125470 | SAMN26179372 |
| 269.B | Kenya | -1.2921 | 36.8219 | SRR18125469 | SAMN26179373 |
| 27.A | Uzbekistan | 39.7681 | 64.4556 | SRR18125468 | SAMN26179374 |
| 270.A | Kenya | -1.2921 | 36.8219 | SRR18125467 | SAMN26179375 |
| 271.A | South Korea | 35.8987 | 127.0392 | SRR18125466 | SAMN26179376 |
| 272.A | Australia | NA | NA | SRR18125464 | SAMN26179377 |
| 273.A | Kenya | -1.2921 | 36.8219 | SRR18125463 | SAMN26179378 |
| 274.A | Kenya | -1.2921 | 36.8219 | SRR18125462 | SAMN26179379 |
| 275.A | Kenya | -1.2921 | 36.8219 | SRR18125461 | SAMN26179380 |
| 276.A | Kenya | -1.2921 | 36.8219 | SRR18125460 | SAMN26179381 |
| 277.B | Kazakhstan | 43.1521 | 68.2581 | SRR18125459 | SAMN26179382 |
| 278.B | Kazakhstan | 43.0631 | 69.0851 | SRR18125458 | SAMN26179383 |
| 279.A | Kazakhstan | 41.5295 | 69.4133 | SRR18125457 | SAMN26179384 |
| 28.A | Russia | 42.1432 | 47.095 | SRR18125456 | SAMN26179385 |
| 280.B | Kazakhstan | 41.5295 | 69.4133 | SRR18125455 | SAMN26179386 |
| 281.B | South Korea | 35.8987 | 127.0392 | SRR18125453 | SAMN26179387 |
| 282.B | South Korea | 35.8987 | 127.0392 | SRR18125452 | SAMN26179388 |
| 283.B | South Korea | 35.8987 | 127.0392 | SRR18125451 | SAMN26179389 |
| 284.A | Tajikistan | 38.0116 | 71.003 | SRR18125450 | SAMN26179390 |
| 285.A | Tajikistan | 37.074793 | 67.957222 | SRR18125449 | SAMN26179391 |
| 286.A | Tajikistan | 37.028926 | 68.004059 | SRR18125448 | SAMN26179392 |
| 287.A | Tajikistan | 39.179338 | 68.012339 | SRR18125447 | SAMN26179393 |
| 288.A | Philippines | 14.5995 | 120.9842 | SRR18125446 | SAMN26179394 |
| 289.B | Philippines | 14.5995 | 120.9842 | SRR18125445 | SAMN26179395 |
| 290.A | China | 40.2374 | 116.2305 | SRR18125444 | SAMN26179396 |
| 290.B | China | 40.2374 | 116.2305 | SRR18125263 | SAMN26179397 |
| 291.A | China | 40.2374 | 116.2305 | SRR18125262 | SAMN26179398 |
| 292.A | China | 30.7378 | 112.2384 | SRR18125261 | SAMN26179399 |
| 293.A | China | 30.7378 | 112.2384 | SRR18125260 | SAMN26179400 |
| 294.A | China | 40.2374 | 116.2305 | SRR18125259 | SAMN26179401 |
| 295.B | China | 37.8957 | 114.9042 | SRR18125258 | SAMN26179402 |
| 296.A | China | 37.8957 | 114.9042 | SRR18125257 | SAMN26179403 |
| 297.A | China | 47.1216 | 128.7382 | SRR18125256 | SAMN26179404 |
| 298.B | China | 47.1216 | 128.7382 | SRR18125255 | SAMN26179405 |
| 299.A | China | 40.2374 | 116.2305 | SRR18125254 | SAMN26179406 |
| 2A | China | 47.1216 | 128.7382 | SRR18125252 | SAMN26179407 |
| 3.B | China | 47.1216 | 128.7382 | SRR18125251 | SAMN26179408 |
| 30.A | Russia | 42.1432 | 47.095 | SRR18125250 | SAMN26179409 |
| 300.A | China | 47.1216 | 128.7382 | SRR18125249 | SAMN26179410 |
| 31.A | Russia | 42.1432 | 47.095 | SRR18125248 | SAMN26179411 |
| 32.B | Russia | 42.1432 | 47.095 | SRR18125247 | SAMN26179412 |
| 33.A | USA | NA | NA | SRR18125246 | SAMN26179413 |
| 34.A | USA | NA | NA | SRR18125245 | SAMN26179414 |
| 34.B | USA | NA | NA | SRR18125244 | SAMN26179415 |
| 35.A | Russia | NA | NA | SRR18125243 | SAMN26179416 |
| 36.A | Russia | 43.1198 | 131.8869 | SRR18125241 | SAMN26179417 |
| 37.A | USA | NA | NA | SRR18125240 | SAMN26179418 |
| 38.B | China | 44.9188 | 130.5244 | SRR18125239 | SAMN26179419 |
| 39.A | Iran | 36.3394 | 59.4698 | SRR18125238 | SAMN26179420 |
| 3A | China | 47.1216 | 128.7382 | SRR18125237 | SAMN26179421 |
| 40.B | Iran | 36.3394 | 59.4698 | SRR18125236 | SAMN26179422 |
| 41.B | Iran | 32.4279 | 53.688 | SRR18125235 | SAMN26179423 |
| 42.A | Iran | 32.4279 | 53.688 | SRR18125234 | SAMN26179424 |
| 43.A | Iran | 32.4279 | 53.688 | SRR18125233 | SAMN26179425 |
| 44.B | Turkmenistan | 37.9153 | 58.0897 | SRR18125232 | SAMN26179426 |
| 45.A | Turkmenistan | 37.9153 | 58.0897 | SRR18125230 | SAMN26179427 |
| 46.A | Turkmenistan | 37.9153 | 58.0897 | SRR18125443 | SAMN26179428 |
| 47.A | Turkmenistan | 37.9153 | 58.0897 | SRR18125442 | SAMN26179429 |
| 48.B | Turkmenistan | 37.9153 | 58.0897 | SRR18125441 | SAMN26179430 |
| 49.B | Turkmenistan | 37.9172 | 58.0907 | SRR18125440 | SAMN26179431 |
| 4A | China | 47.1216 | 128.7382 | SRR18125439 | SAMN26179432 |
| 5.B | China | 47.1216 | 128.7382 | SRR18125438 | SAMN26179433 |
| 50.A | Turkmenistan | 37.9172 | 58.0907 | SRR18125437 | SAMN26179434 |
| 51.B | Turkmenistan | 37.9601 | 58.3261 | SRR18125436 | SAMN26179435 |
| 52.A | Turkmenistan | 37.9601 | 58.3261 | SRR18125435 | SAMN26179436 |
| 53.B | USA | 40.1605 | -103.2144 | SRR18125433 | SAMN26179437 |
| 54.A | USA | 40.1605 | -103.2144 | SRR18125432 | SAMN26179438 |
| 55.B | Ukraine | 48.3794 | 31.1656 | SRR18125431 | SAMN26179439 |
| 56.A | Kazakhstan | 43.3667 | 68.4094 | SRR18125430 | SAMN26179440 |
| 57.A | Iran | 34.7608 | 48.3988 | SRR18125429 | SAMN26179441 |
| 58.A | Iran | 35.6892 | 51.389 | SRR18125428 | SAMN26179442 |
| 59.B | Kazakhstan | 43.3667 | 68.4094 | SRR18125427 | SAMN26179443 |
| 6.B | China | 47.1216 | 128.7382 | SRR18125426 | SAMN26179444 |
| 60.A | Kazakhstan | 42.2663 | 68.1431 | SRR18125425 | SAMN26179445 |
| 61.A | Uzbekistan | 41.2995 | 69.2401 | SRR18125424 | SAMN26179446 |
| 62.A | Uzbekistan | 41.2995 | 69.2401 | SRR18125422 | SAMN26179447 |
| 63.A | Uzbekistan | 41.2995 | 69.2401 | SRR18125421 | SAMN26179448 |
| 64.B | Uzbekistan | 41.2995 | 69.2401 | SRR18125420 | SAMN26179449 |
| 65.B | Uzbekistan | 41.2995 | 69.2401 | SRR18125419 | SAMN26179450 |
| 66.B | Uzbekistan | 41.2995 | 69.2401 | SRR18125418 | SAMN26179451 |
| 67.A | Uzbekistan | 41.2995 | 69.2401 | SRR18125417 | SAMN26179452 |
| 68.A | Uzbekistan | 41.2995 | 69.2401 | SRR18125416 | SAMN26179453 |
| 69.A | Uzbekistan | 40.4915 | 68.7811 | SRR18125415 | SAMN26179454 |
| 70.A | Uzbekistan | 39.627 | 66.975 | SRR18125414 | SAMN26179455 |
| 71.A | Uzbekistan | 39.627 | 66.975 | SRR18125413 | SAMN26179456 |
| 72.A | Uzbekistan | 39.627 | 66.975 | SRR18125411 | SAMN26179457 |
| 73.B | Uzbekistan | 39.627 | 66.975 | SRR18125410 | SAMN26179458 |
| 74.A | Uzbekistan | 39.627 | 66.975 | SRR18125409 | SAMN26179459 |
| 75.A | Uzbekistan | 39.627 | 66.975 | SRR18125408 | SAMN26179460 |
| 76.A | Uzbekistan | 41.2995 | 69.2401 | SRR18125229 | SAMN26179461 |
| 77.A | Uzbekistan | 39.7681 | 64.4556 | SRR18125228 | SAMN26179462 |
| 79.B | Uzbekistan | 39.7681 | 64.4556 | SRR18125227 | SAMN26179463 |
| 7A | China | 47.1216 | 128.7382 | SRR18125226 | SAMN26179464 |
| 8.B | China | 47.1216 | 128.7382 | SRR18125225 | SAMN26179465 |
| 81.B | Afghanistan | 34.1769 | 61.7006 | SRR18125224 | SAMN26179466 |
| 82.B | Afghanistan | 34.1769 | 61.7006 | SRR18125222 | SAMN26179467 |
| 83.A | Afghanistan | 34.1769 | 61.7006 | SRR18125221 | SAMN26179468 |
| 84.A | Afghanistan | 34.1769 | 61.7006 | SRR18125220 | SAMN26179469 |
| 85.A | Afghanistan | 34.1769 | 61.7006 | SRR18125219 | SAMN26179470 |
| 86.A | Afghanistan | 36.6153 | 66.9293 | SRR18125218 | SAMN26179471 |
| 88.B | Afghanistan | NA | NA | SRR18125217 | SAMN26179472 |
| 89.B | Afghanistan | 33.9391 | 67.71 | SRR18125216 | SAMN26179473 |
| 90.B | Uzbekistan | NA | NA | SRR18125215 | SAMN26179474 |
| 91.A | Japan | 35.719 | 139.7456 | SRR18125214 | SAMN26179475 |
| 92.A | Japan | 35.719 | 139.7456 | SRR18125213 | SAMN26179476 |
| 93.B | Armenia | 40.0691 | 45.0382 | SRR18125211 | SAMN26179477 |
| 94.A | Iran | 35.102 | 59.1042 | SRR18125210 | SAMN26179478 |
| 95.A | Azerbaijan | 40.4093 | 49.8671 | SRR18125209 | SAMN26179479 |
| 96.A | Azerbaijan | 40.1431 | 47.5769 | SRR18125208 | SAMN26179480 |
| 97.A | Turkmenistan | 39.0041 | 63.5688 | SRR18125207 | SAMN26179481 |
| 98.B | Panama | 9.3593 | -79.8999 | SRR18125206 | SAMN26179482 |
| 99.A | Panama | 9.3593 | -79.8999 | SRR18125205 | SAMN26179483 |
| 99.B | Panama | 9.3593 | -79.8999 | SRR18125204 | SAMN26179484 |
| 9A | China | 47.1216 | 128.7382 | SRR18125203 | SAMN26179485 |
| M7.A | India | 31.5204 | 74.3587 | SRR18125202 | SAMN26179486 |

**Supplementary file 2.** Outgroup *f3* statistics among all possible combinations of genetic group pmrs

| Outgroup<br>(C) | Source<br>1<br>(A) | Source2<br>(B) | $f_3$ | Standard error | Z-score | Significant |
| --- | --- | --- | --- | --- | --- | --- |
| *sublobata* | CA | EA | 0.232 | 0.005 | 42.66 | Yes |
| *sublobata* | CA | SA | 0.209 | 0.005 | 38.41 | Yes |
| *sublobata* | CA | SEA | 0.213 | 0.005 | 39.03 | Yes |
| *sublobata* | EA | SA | 0.209 | 0.005 | 38.79 | Yes |
| *sublobata* | EA | SEA | 0.218 | 0.005 | 40.75 | Yes |
| *sublobata* | SA | SEA | 0.211 | 0.005 | 39.41 | Yes |

Abbreviations: SA, South Asia; SEA, Southeast Asia; EA, East Asia and CA, Central Asia ( $f_3$ statistics with Z-score $> |3|$ are considered significant)

**Supplementary file 3.** Admixture *f3* statistics among all possible population trios

| Target<br>(C) | Source1<br>(A) | Source2<br>(B) | $f_3$ | Standard error | Z-score | Significant |
| --- | --- | --- | --- | --- | --- | --- |
| EA | SA | CA | 0.005 | 0.001 | 4.82 | Yes |
| EA | SEA | CA | -0.001 | 0.001 | -0.51 | No |
| EA | SEA | SA | 0.020 | 0.002 | 13.48 | Yes |
| SEA | CA | EA | 0.030 | 0.002 | 14.7 | Yes |
| SEA | SA | CA | 0.014 | 0.002 | 8.82 | Yes |
| SEA | SA | EA | 0.009 | 0.001 | 6.86 | Yes |
| SA | CA | EA | 0.032 | 0.002 | 16.66 | Yes |
| SA | CA | SEA | 0.011 | 0.001 | 9.02 | Yes |
| SA | EA | SEA | 0.017 | 0.001 | 12.91 | Yes |
| CA | EA | SA | 0.011 | 0.001 | 9.55 | Yes |
| CA | EA | SEA | 0.016 | 0.002 | 10.37 | Yes |
| CA | SEA | SA | 0.031 | 0.002 | 15.65 | Yes |

Abbreviations: SA, South Asia; SEA, Southeast Asia; EA, East Asia and CA, Central Asia ( $f_3$ statistics with Z-score $> |3|$ are considered significant, but only negative $f_3$ statistics denote the target population being admixed from source1 and source2.)

**Supplementary file 4.** Mantel tests for isolation by distance of inferred genetic group (Q 2 0.5)

| **Group** | ***r*** | ***P*** |
| --- | --- | --- |
| SA | 0.4319 | 0.008* |
| SEA | 0.3312 | 0.041* |
| EA | 0.0461 | 0.052 |
| CA | 0.0070 | 0.435 |
| Southern | 0.2934 | 0.001* |
| Northern | 0.2777 | 0.001* |

Abbreviations: SA, South Asia; SEA, Southeast Asia; EA, East Asia; CA, Central Asia; *r*, Mantel correlation; significance level \* $P < 0.05$

**Supplementary file 5.**
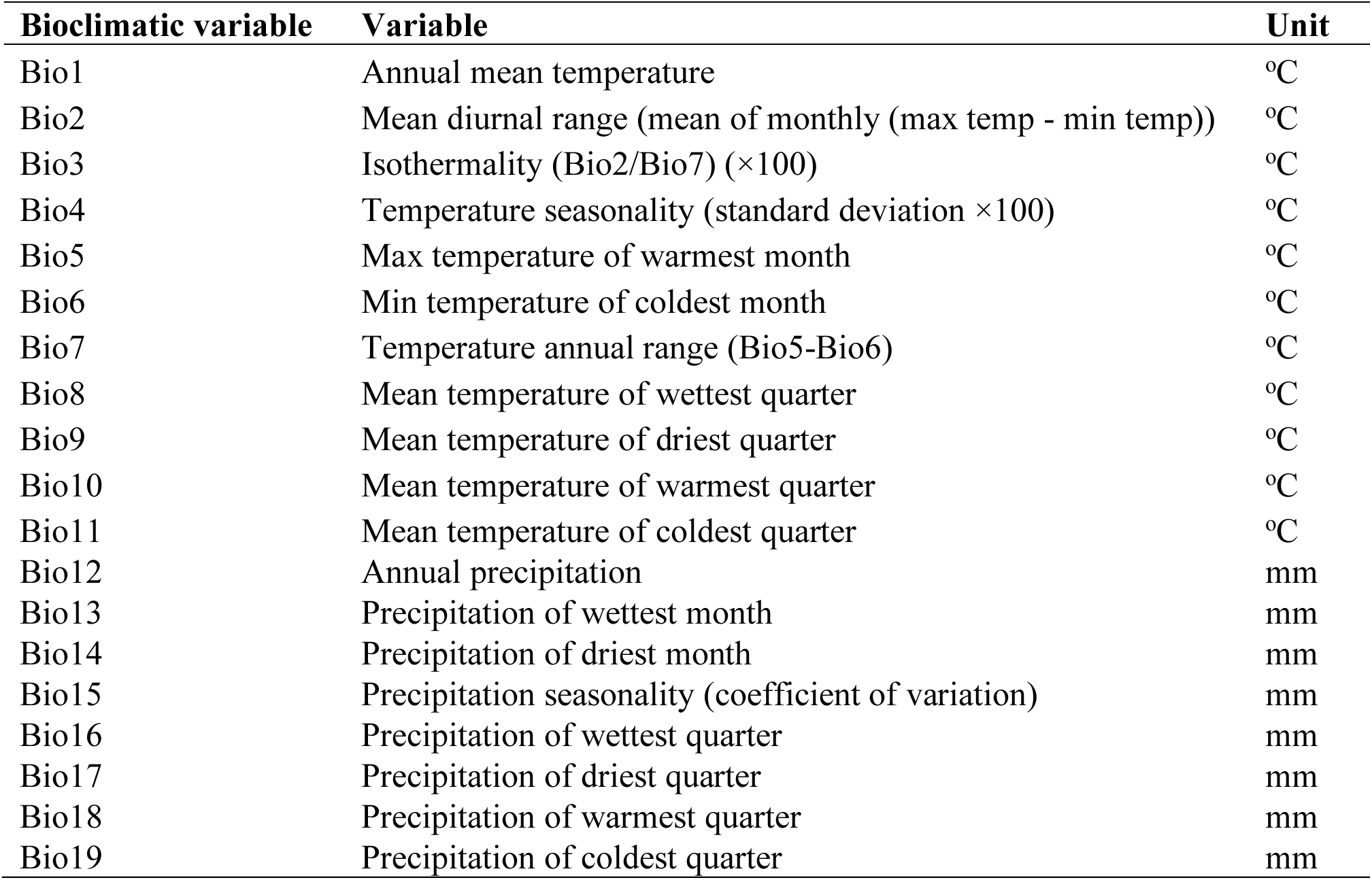
Description of bioclimatic variables used in ecological niche modelling

**Supplementary file 6.** Pearson’s correlation coefficient between pairs of bioclimatic variables (denoted in lower triangle)

| **Bioclimatic variable** | **Bio1** | **Bio2** | **Bio3** | **Bio4** | **Bio5** | **Bio6** | **Bio7** | **Bio8** | **Bio9** | **Bio10** | **Bio11** | **Bio12** | **Bio13** | **Bio14** | **Bio15** | **Bio16** | **Bio17** | **Bio18** | **Bio19** |
| --- | --- | --- | --- | --- | --- | --- | --- | --- | --- | --- | --- | --- | --- | --- | --- | --- | --- | --- | --- |
| **Bio1** | 1 | - | - | - | - | - | - | - | - | - | - | - | - | - | - | - | - | - | - |
| **Bio2** | 0.087 | 1 | - | - | - | - | - | - | - | - | - | - | - | - | - | - | - | - | - |
| **Bio3** | 0.732 | 0.321 | 1 | - | - | - | - | - | - | - | - | - | - | - | - | - | - | - | - |
| **Bio4** | -0.814 | 0.011 | -0.876 | 1 | - | - | - | - | - | - | - | - | - | - | - | - | - | - | - |
| **Bio5** | 0.872 | 0.258 | 0.455 | -0.449 | 1 | - | - | - | - | - | - | - | - | - | - | - | - | - | - |
| **Bio6** | 0.971 | -0.038 | 0.779 | -0.911 | 0.749 | 1 | - | - | - | - | - | - | - | - | - | - | - | - | - |
| **Bio7** | -0.773 | 0.225 | -0.784 | 0.973 | -0.377 | -0.896 | 1 | - | - | - | - | - | - | - | - | - | - | - | - |
| **Bio8** | 0.595 | -0.066 | 0.383 | -0.398 | 0.522 | 0.530 | -0.392 | 1 | - | - | - | - | - | - | - | - | - | - | - |
| **Bio9** | 0.901 | 0.122 | 0.663 | -0.759 | 0.803 | 0.896 | -0.715 | 0.257 | 1 | - | - | - | - | - | - | - | - | - | - |
| **Bio10** | 0.915 | 0.129 | 0.472 | -0.513 | 0.983 | 0.808 | -0.472 | 0.590 | 0.815 | 1 | - | - | - | - | - | - | - | - | - |
| **Bio11** | 0.977 | 0.050 | 0.814 | -0.918 | 0.760 | 0.994 | -0.881 | 0.544 | 0.894 | 0.811 | 1 | - | - | - | - | - | - | - | - |
| **Bio12** | 0.237 | -0.513 | 0.244 | -0.397 | -0.030 | 0.327 | -0.477 | 0.293 | 0.121 | 0.073 | 0.303 | 1 | - | - | - | - | - | - | - |
| **Bio13** | 0.276 | -0.341 | 0.289 | -0.403 | 0.040 | 0.332 | -0.438 | 0.342 | 0.142 | 0.124 | 0.330 | 0.926 | 1 | - | - | - | - | - | - |
| **Bio14** | -0.084 | -0.620 | -0.151 | -0.011 | -0.205 | 0.013 | -0.154 | 0.039 | -0.120 | -0.126 | -0.048 | 0.494 | 0.233 | 1 | - | - | - | - | - |
| **Bio15** | 0.258 | 0.528 | 0.405 | -0.263 | 0.208 | 0.191 | -0.128 | 0.302 | 0.156 | 0.189 | 0.262 | -0.024 | 0.213 | -0.521 | 1 | - | - | - | - |
| **Bio16** | 0.271 | -0.364 | 0.283 | -0.406 | 0.028 | 0.333 | -0.446 | 0.330 | 0.141 | 0.115 | 0.328 | 0.950 | 0.992 | 0.258 | 0.178 | 1 | - | - | - |
| **Bio17** | -0.050 | -0.639 | -0.110 | -0.058 | -0.194 | 0.052 | -0.203 | 0.062 | -0.089 | -0.110 | -0.009 | 0.555 | 0.294 | 0.988 | -0.508 | 0.321 | 1 | - | - |
| **Bio18** | -0.031 | -0.453 | -0.011 | -0.119 | -0.257 | 0.027 | -0.210 | 0.245 | -0.184 | -0.144 | 0.015 | 0.805 | 0.741 | 0.392 | 0.016 | 0.766 | 0.438 | 1 | - |
| **Bio19** | 0.177 | -0.270 | 0.266 | -0.279 | 0.048 | 0.263 | -0.335 | -0.010 | 0.218 | 0.080 | 0.227 | 0.460 | 0.353 | 0.468 | -0.196 | 0.356 | 0.490 | 0.115 | 1 |

**Supplementary file 7.**
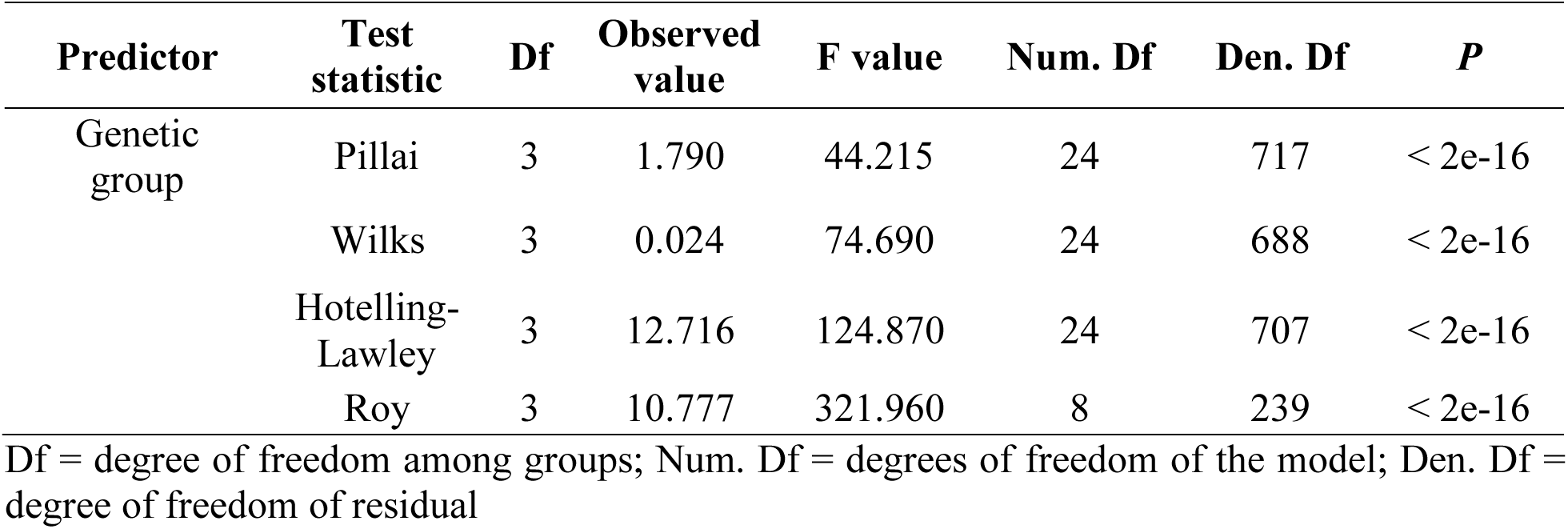
Comparison of bioclimatic variables among the four genetic groups analysed with multivariate analysis of variance (MANOVA)

**Supplementary file 8.** Summary of analysis of variance (ANOVA) for bioclimatic variables

Df = degree of freedom among groups; Num. Df = degrees of freedom of the model; Den. Df = degree of freedom of residual **Supplementary file 8.** Summary of analysis of variance (ANOVA) for bioclimatic variables
| **Bioclimatic variable** | **Df** | **Sum square** | **Mean square** | **F value** | ***P*** |
| --- | --- | --- | --- | --- | --- |
| Bio1 (Annual temperature) | 3 | 183.688 | 61.229 | 235.97 | <2.2e-16 |
| Bio2 (Mean diurnal temperature range) | 3 | 66.253 | 22.085 | 29.813 | <2.2e-16 |
| Bio3 (Isothermality) | 3 | 188.031 | 62.677 | 259.34 | <2.2e-16 |
| Bio8 (Mean temperature of wettest quarter) | 3 | 177.063 | 59.021 | 205.91 | <2.2e-16 |
| Bio12 (Annual precipitation) | 3 | 157.890 | 52.630 | 144.11 | <2.2e-16 |
| Bio14 (Precipitation of driest month) | 3 | 29.859 | 9.952 | 11.183 | <6.6e-07 |
| Bio15 (Precipitation seasonality) | 3 | 119.810 | 39.938 | 76.62 | <2.2e-16 |
| Bio19 (Precipitation of coldest quarter) | 3 | 44.212 | 14.737 | 17.732 | < 1.9e-10 |

Df = degree of freedom

**Supplementary file 9.** Correlation between eight bioclimatic variables and climatic PC axes 1 to 4

Df = degree of freedom **Supplementary file 9.** Correlation between eight bioclimatic variables and climatic PC axes 1 to 4
| Bioclimatic variable | PC1 | PC2 | PC3 | PC4 |
| --- | --- | --- | --- | --- |
| Bio1 (Annual temperature) | **-0.445** | -0.107 | 0.390 | -0.056 |
| Bio2 (Mean diurnal temperature range) | 0.193 | **-0.564** | 0.062 | 0.253 |
| Bio3 (Isothermality) | **-0.430** | -0.082 | **0.492** | -0.184 |
| Bio8 (Mean temperature of wettest quarter) | **-0.486** | 0.042 | -0.287 | 0.049 |
| Bio12 (Annual precipitation) | **-0.401** | 0.366 | 0.066 | 0.202 |
| Bio14 (Precipitation of driest month) | 0.017 | **0.582** | -0.248 | 0.259 |
| Bio15 (Precipitation seasonality) | -0.323 | -0.365 | -0.292 | **0.667** |
| Bio19 (Precipitation of coldest quarter) | 0.279 | 0.235 | **0.607** | **0.587** |

(Correlation coefficients with absolute values higher than 0.4 are in bold.)

**Supplementary file 10.** Comparison of summer growing season data including temperature and precipitation of May, July and September among the four genetic groups analysed with multivariate analysis of variance (MANOVA)

| Predictor | Test statistic | Df | Observed value | F value | Num. Df | Den. Df | *P* |
| --- | --- | --- | --- | --- | --- | --- | --- |
| Genetic group | Pillai | 3 | 1.915 | 70.907 | 18 | 723 | < 2e-16 |
|  | Wilks | 3 | 0.010 | 156.350 | 18 | 676 | < 2e-16 |
|  | Hotelling-Lawley | 3 | 19.873 | 262.400 | 18 | 713 | < 2e-16 |
|  | Roy | 3 | 15.400 | 618.570 | 6 | 241 | < 2e-16 |

Df = degree of freedom among groups; Num. Df = degrees of freedom of the model; Den. Df = degree of freedom of residual

**Supplementary file 11.** ANOVA table for all evaluated field traits (phenology, reproduction and size) as well as drought-related traits

1095
| Trait | Garden | Model $r^2$ | Group $F$ | Group $P$ | SEA<sup>1</sup> | SA<sup>1</sup> | CA<sup>1</sup> | Tukey<sup>2</sup> |
| --- | --- | --- | --- | --- | --- | --- | --- | --- |
| **Phenology:** |  |  |  |  |  |  |  |  |
| Days to 50% flowering | Pakistan 2015 | 0.2388 | 7.2144 | 0.0019* | -0.3702 | 0.6296 | -0.3525 | B,A,B |
| Days to 50% flowering | Taiwan 1984 | 0.5266 | 25.5887 | <.0001* | 0.5723 | 0.3197 | -1.0685 | A,A,B |
| Days to 50% flowering | Taiwan 2018 | 0.4465 | 18.5544 | <.0001* | 0.0974 | 0.6569 | -0.9633 | A,A,B |
| **Reproduction:** |  |  |  |  |  |  |  |  |
| 100 seed weight, g | Pakistan 2015 | 0.4722 | 20.5761 | <.0001* | 0.9050 | -0.6397 | -0.2756 | A,B,B |
| Pod length, cm | Pakistan 2015 | 0.2621 | 8.1698 | 0.0009* | 0.6810 | -0.2605 | -0.4896 | A,B,B |
| Pod length, cm | Taiwan 1984 | 0.7173 | 58.3615 | <.0001* | 1.0524 | -0.1980 | -1.0232 | A,B,C |
| Pods per plant | Pakistan 2015 | 0.4471 | 18.5988 | <.0001* | -0.6441 | 0.8455 | -0.3062 | B,A,B |
| 1000 seed weight, g | Taiwan 1984 | 0.6248 | 38.3053 | <.0001* | 0.9979 | -0.8380 | -0.1340 | A,C,B |
| Seed yield per plant, g | Pakistan 2015 | 0.4666 | 20.1225 | <.0001* | -0.6210 | 0.8752 | -0.3712 | B,A,B |
| Seeds per pod | Pakistan 2015 | 0.1300 | 3.4372 | 0.0406* | -0.4875 | 0.2418 | 0.2806 | A,A,A |
| Seeds per pod | Taiwan 1984 | 0.1611 | 4.4168 | 0.0176* | 0.1413 | 0.3386 | -0.6107 | AB,A,B |
| **Plant size:** |  |  |  |  |  |  |  |  |
| Petiole length, cm | Pakistan 2015 | 0.2943 | 9.5907 | 0.0003* | 0.5435 | 0.0878 | -0.7798 | A,A,B |
| Plant height, cm | Pakistan 2015 | 0.0001 | 0.0024 | 0.9976 | 0.0075 | 0.0055 | -0.0158 | A,A,A |
| Plant height at flowering, cm | Taiwan 1984 | 0.3981 | 15.2115 | <.0001* | 0.4811 | 0.3024 | -0.9705 | A,A,B |
| Plant height at maturity, cm | Taiwan 1984 | 0.5472 | 27.8000 | <.0001* | 0.3480 | 0.5605 | -1.1362 | A,A,B |
| Primary leaf length, cm | Taiwan 1984 | 0.5454 | 27.5930 | <.0001* | 0.9813 | -0.4212 | -0.6253 | A,B,B |
| Primary leaf width, cm | Taiwan 1984 | 0.6053 | 35.2773 | <.0001* | 1.0244 | -0.6010 | -0.4313 | A,B,B |
| Terminal leaflet length, cm | Pakistan 2015 | 0.2186 | 6.4340 | 0.0034* | 0.3062 | 0.2643 | -0.7167 | A,A,B |
| Terminal leaflet width, cm | Pakistan 2015 | 0.1680 | 4.6458 | 0.0145* | 0.4361 | 0.0387 | -0.5734 | A,AB,B |
| **Drought (PEG6000):** |  |  |  |  |  |  |  |  |
| Shoot dry weight (SDW), mg | NTU 2021 | 0.5998 | 36.7246 | <.0001* | 1.0508 | -0.5607 | -0.5483 | A,B,B |
| Root dry weight (RDW), mg | NTU 2021 | 0.5964 | 36.2048 | <.0001* | 1.0299 | -0.6933 | -0.3336 | A,B,B |
| Total dry weight (TDW), mg | NTU 2021 | 0.5934 | 35.7555 | <.0001* | 1.0448 | -0.5883 | -0.5028 | A,B,B |
| Root:Shoot ratio dry weight (RSRDW) | NTU 2021 | 0.3261 | 11.8577 | <.0001* | -0.2316 | -0.4342 | 0.9112 | B,B,A |
| **Drought (Control):** |  |  |  |  |  |  |  |  |
| Shoot dry weight (SDW), mg | NTU 2021 | 0.5779 | 33.5453 | <.0001* | 1.0275 | -0.4709 | -0.6484 | A,B,B |
| Root dry weight (RDW), mg | NTU 2021 | 0.5205 | 26.5962 | <.0001* | 0.9716 | -0.6077 | -0.3811 | A,B,B |
| Total dry weight (TDW), mg | NTU 2021 | 0.5638 | 31.6722 | <.0001* | 1.0195 | -0.5374 | -0.5430 | A,B,B |
| Root:Shoot ratio dry weight (RSRDW) | NTU 2021 | 0.1753 | 5.2083 | 0.0089* | -0.3036 | -0.2004 | 0.6773 | B,B,A |

\*Significant at $P < 0.05$ ; 1: least-square means of each group after inverse normal transformation of raw data; 2: levels not connected by same letter are significantly different.

**Supplementary file 12.** Mean of eight bioclimatic variables of the genetic groups

| **Bioclimatic**<br>**variable** | **Northeast Asia**<br>**(N = 37)**<br>**Mean ± SD** | **Northwest Asia**<br>**(N = 45)**<br>**Mean ± SD** | **Southeast Asia**<br>**(N = 45)**<br>**Mean ± SD** | **South Asia**<br>**(N = 49)**<br>**Mean ± SD** | **Central Asia**<br>**(N = 72)**<br>**Mean ± SD** |
| --- | --- | --- | --- | --- | --- |
| **Bio1** | 62.49 ± 55.39 | 117.58 ± 54.30 | 256.47 ± 19.29 | 255.57 ± 14.80 | 128.72 ± 41.10 |
| **Bio2** | 115.22 ± 15.44 | 124.38 ± 20.14 | 102.02 ± 16.11 | 127.24 ± 16.12 | 130.08 ± 13.89 |
| **Bio3** | 25.14 ± 2.04 | 31.02 ± 4.47 | 51.58 ± 7.19 | 42.8 ± 4.58 | 32.4 ± 3.52 |
| **Bio8** | 207.86 ± 31.97 | 107.89 ± 67.90 | 271.47 ± 11.68 | 283.92 ± 23.02 | 92.04 ± 42.23 |
| **Bio12** | 821.59 ± 299.98 | 301.24 ± 201.00 | 1477.69 ± 380.18 | 750.39 ± 329.19 | 285.67 ± 145.24 |
| **Bio14** | 12.14 ± 11.72 | 5.18 ± 8.99 | 6.33 ± 4.34 | 2.47 ± 3.44 | 2.51 ± 4.98 |
| **Bio15** | 95.35 ± 24.52 | 65.4 ± 20.35 | 84.78 ± 7.25 | 124.47 ± 23.48 | 70.92 ± 12.92 |
| **Bio19** | 42.51 ± 38.23 | 88.33 ± 67.16 | 48.02 ± 24.91 | 34 ± 33.99 | 92.15 ± 45.19 |

